# Plant genotype and microbial strain combinations strongly influence the transcriptome under heavy metal stress conditions

**DOI:** 10.1101/2024.10.06.616885

**Authors:** Reena Sharma, Sanhita Chakraborty, Aditi Bhat, Michael Clear, Meng Xie, José J. Pueyo, Timothy Paape

## Abstract

Heavy metals such as cadmium (Cd) and mercury (Hg) pose significant threats to plant health and food safety as they are absorbed from the environment. Legumes are generally considered sensitive to heavy metals but possess standing genetic variation for accumulation and tolerance to toxic ions. We conducted a transcriptomic analysis on hydroponically and soil grown *Medicago truncatula* plants to investigate gene expression responses to Cd and Hg exposure in roots, leaves, and nodules. By using plant genotypes with varying metal tolerance or accumulation levels, we observed distinct clustering of gene ontologies, indicating tissue-specific, genotype-specific, and metal-specific gene expression patterns. Considering the symbiotic relationship between legumes and nitrogen-fixing bacteria, we further examined plant phenotypes and transcriptomes of plant genotypes with contrasting Hg accumulation levels and inoculated them with high or low Hg-tolerant *Sinorhizobium medicae* strains that have presence-absence variation for a mercury reductase (Mer) operon. Host plants inoculated with the Hg-tolerant rhizobia strain possessing a Mer operon exhibited less reduction in nodule number and plant biomass. A smaller reduction in iron (Fe) distribution in nodules after Hg stress was measured using X-ray Fluorescence (XRF) imaging. Dual transcriptome (host plant and bacteria) analysis of nodules revealed a remarkable decrease in the number of differentially expressed genes (DEGs) and clustering of gene ontologies in plants inoculated with the Hg-tolerant rhizobia strain, including symbiosis related genes. This suggests the Hg-tolerant rhizobia strain has the potential to mitigate Hg stress in host plants. Furthermore, we observed genotype-by-genotype interactions between the high Hg accumulating plant genotype and the Hg-tolerant rhizobia strain. These findings provide insights into enhancing plant resilience in contaminated environments through optimizing legume-rhizobia interactions for heavy metal tolerance, as well as identification of genetic mechanisms that can reduce transport to edible plant parts.

**Highlights:**

- Genetic variation in tolerance to heavy metals affects responses in plant tissues
- Symbiosis with Hg tolerant rhizobia strain reduced stress response in host plant
- Dual-transcriptomics in nodules identified metal stress and symbiosis responses
- Genotype by genotype interaction significantly affected symbiosis gene expression
- Rhizobia strain possessing Mer operon suggests genetic basis affecting responses

## 1. Introduction

Increased anthropogenic activities such as the construction of mines, foundries, and smelters, and the excessive application of pesticides and herbicides create marginal agricultural soils with accumulation of heavy metals such as cadmium (Cd) and mercury (Hg), leading to phytotoxicity and microbial toxicity (Alloway, 2013; Tchounwou et al., 2012). These metals can accumulate in the aerial parts of plants, including leaves and seeds, and enter into the food supply (Alengebawy et al., 2021; Peralta-Videa et al., 2009; Rai et al., 2019). Moreover, these toxic heavy metals pose a considerable risk to ecological and agricultural systems by disrupting microbiomes and plant health (Sachdev and Ansari, 2022; Singh et al., 2023).

Plants use a complex array of detoxification mechanisms to combat the toxicity of Cd and Hg. Heavy metals primarily enter plants through root absorption via transporters during import of essential nutrients, and impact metabolism and micronutrient homeostasis through the increased production of reactive oxygen species (ROS) and oxidative stress (Mansoor et al., 2023). Therefore, plant roots are the first line of defense to protect aerial parts of the plants, and tend to show the largest stress responses in the transcriptome. Glutathione S-transferases (GSTs) play a crucial role in the detoxification of Cd and Hg in plants through the activation of the phytochelatin synthase (PCS) enzymatic pathway (Cobbett, 2001; Hossain et al., 2012), leading to chelation and can be transported into plant vacuoles. GSTs catalyze the conjugation of reduced glutathione to these heavy metal ions (Kumar and Trivedi, 2018), followed by compartmentalization of Cd and Hg in roots and leaves by chelation of these metals using phytochelatins and metallothioneins (Clemens, 2006) which bind to Cd and Hg to form more stable, less toxic complexes (Yadav, 2010). These complexes are then actively transported using ATP-binding cassette (ABC) transporters for sequestration or expulsion from the cell (Kang et al., 2011; Park et al., 2012), reducing their harmful concentrations in the cytosol. Additionally, unchelated metal ions can be directly sequestered into the vacuole using cation diffusion facilitators such as ABCs, ATPases, and MTPs (metal tolerance proteins) (El-Sappah et al., 2021; León-Mediavilla et al., 2018; Shahzad et al., 2010).

Plants exposed to heavy metals trigger the production of excess ROS as a result of disruption of mitochondrial membranes and impairment of enzymes that can clear ROS in the cellular antioxidant process. Thus, accumulation of ROS can result in oxidative stress, which damages various cellular components including proteins, lipids, and DNA. The oxidative damage then disrupts cellular functions, leading to impaired growth, reduced photosynthesis, and ultimately cell death. The oxidative stress induced by these metals can be mitigated by the upregulation of antioxidant enzymes like superoxide dismutase (SOD), catalase (CAT), lipoxygenase (LOX) and peroxidases (Kryvoruchko et al., 2018; Sachdev et al., 2021). This integrated response, encompassing chelation, sequestration, transport, and antioxidant pathways, is crucial for plant survival in environments contaminated with heavy metals.

The model legume *Medicago truncatula* is widely distributed across diverse geographical regions, very often in dry soils with high salinity and marginal conditions (Yoder et al., 2014). Some populations of *Medicago* sp. have been found in contaminated mine sites with high levels of Cd, Hg, and Zn (García de la Torre et al., 2013; Nonnoi et al., 2012). *Medicago* species form symbiosis with nitrogen-fixing soil rhizobia, which produces nodules on the host plant roots that contain a mixture of plant and bacteria cells. When plants are exposed to heavy metals such as Cd and Hg, cellular homeostasis is disrupted, and legume-rhizobium symbiosis depends on maintaining ion homeostasis in the roots and nodules (Chakraborty and Harris, 2022; Rodríguez-Haas et al., 2013). Oxidative stress induced by heavy metal toxicity generally disrupts the symbiosis (Hawkins and Oresnik, 2022). In response to soil stress, such as salinity, nutrient deficiency, or soil toxicity, legumes may modulate the expression of nodulation and symbiosis related genes. This regulation can disrupt normal nodulation such as reducing the number of nodules formed or altering their distribution on the roots (Escudero et al., 2020a; León-Mediavilla et al., 2018). This helps to optimize resource allocation and minimize the energy cost associated with nodulation during stress conditions. Given that various *Medicago* species, such as *M. sativa* (alfalfa) serve as forage crops, it is essential to prevent the translocation of toxic ions into edible plant parts to avoid their entry into the food chain.

We conducted an experiment using *M. truncatula* seedlings under hydroponic conditions to identify transcriptomic responses in root and leaf tissue of several genotypes from the Medicago HapMap panel which showed variation in leaf accumulation and relative root growth following Cd and Hg stress treatments. There is a relationship between the level of genotypic (or species) resilience to heavy metal stress and their transcriptomic responses, exhibited by fewer differentially expressed genes (DEGs) in resilient genotypes (Bhat et al., 2024; Paape et al., 2016) due to adaptations, while less resilient genotypes often show high numbers of DEGs due to the high impact of the stress to the organism. The hydroponic experimental design follows a recent genome wide association study (GWAS) in *M. truncatula* which revealed the associations of several ABC-transporters, ATPases, and other metal transporters such as MtNRAMP6 with Cd and Hg tolerance and accumulation, which comprised both root and leaf level responses (Paape et al., 2022). While GWAS can identify candidate SNPs in or near genes associated with tolerance and accumulation of Cd and Hg, SNPs alone do not provide information about gene expression responses to the stress. In a second experiment using full grown *M. truncatula* plants inoculated with both low- and high-Hg-tolerant (“LT” and “HT”, respectively) *S. medicae* strains, we induced heavy metal stress in nodules. This approach enabled us to assess the impact of rhizobia genotype on the host plant transcriptome and other plant phenotypes to identify beneficial host plant and rhizobia genotype interactions.

## 2. Materials and Methods

### 2.1 Plant growth using hydroponic conditions for RNA collection from leaf and root tissues

The Medicago HapMap panel was assessed for 4 traits following heavy metal treatments of 10μM CdCl_2_ or 4μM HgCl_2_: relative root growth (RRG) following Hg treatment, RRG following Cd treatment, Cd and Hg accumulation in leaf tissues after the treatment period. The original phenotyping was done on a collection of 230 genotypes that provided the basis for the samples used for the current study. We selected 4 plant genotypes from the Medicago HapMap based phenotype data for accumulation of Cd or Hg and RRG (HM195, HM075, HM302, HM304 and one genotype from the United States Department of Agriculture (USDA) GRIN collection (PI660407) that shows high Cd and Hg accumulation trait (García de la Torre et al., 2013, 2021) (Table S1). To germinate the seeds, they were scarified with sandpaper and sterilized in 50% (v/v) commercial bleach for 15 minutes, then were rinsed 4 times for 20 minutes with sterile water. Seeds were imbibed in sterile water at 4°C overnight and then germinated on 1% agar/water in Petri dishes in growth chamber conditions (25/19°C, 16/8 hours) for 48 hours in darkness. Seeds from *M. truncatula* genotypes were obtained through the Medicago HapMap project (https://medicagohapmap2.org/germplasm). All genotypes were cultured in parallel with two different heavy metal treatments: Cd and Hg. Phenotyping assays were conducted hydroponically in a modified Hoagland nutrient solution based on (García de la Torre et al., 2013) using the following concentrations: 2.02g/L KNO_3_, 0.68g/L KH_2_PO_4_, 0.182g/L CaCl_2_·2H_2_O, 0.615g/L MgSO_4_·7H_2_O, 0.109g/L K_2_SO_4_, 0.205g/L Hampiron (Rhône Poulenc), and 1.35mL of a solution containing: 11g/L H_3_BO_3_, 6.2g/L MnSO_4_·H_2_O, 10g/L KCl, 1g/L ZnSO_4_·7H_2_O, 1g/L (NH_4_)_6_ MO_7_O_2_·4H_2_O, 0.5g/L CuSO_4_·5H_2_O and 0.5mL/L H_2_SO_4_. Cd-treated plants received 10μM CdCl_2_ amended Hoagland solution while Hg-treated plants received 4μM HgCl_2_ amended Hoagland nutrient solution. A control group of plants were cultured in parallel with unamended Hoagland nutrient solution. Three replicate plant seedlings per *M. truncatula* genotype were collected for RNAseq. Seedlings were acclimatized in a growth chamber with 250mL of nutrient solution for 24 hours, followed by 48 hours with or without Cd and Hg treatment. Leaves and roots were harvested after 72 hours plants had been exposed to Cd or Hg treatment. Metal concentration was measured as reported by Paape et al., (2022).

### 2.2 RNA extraction and sequencing libraries of leaf and root samples

The RNA from the hydroponics experiment was extracted using the Qiagen RNAeasy Plant Mini Kit using the standard protocol and sent to the Joint Genome Institute (JGI) for library construction, sequencing, and preliminary analysis of raw reads. Plate-based RNA sample prep was performed on the PerkinElmer Sciclone NGS robotic liquid handling system using Illumina’s TruSeq Stranded mRNA HT sample prep kit utilizing poly-A selection of mRNA following the protocol outlined by Illumina in their user guide: https://support.illumina.com/sequencing/sequencing_kits/truseq-stranded-mrna.html, and with the following conditions: total RNA starting material was 1000ng per sample and 8 cycles of PCR was used for library amplification. The prepared libraries were quantified using KAPA Biosystems next-generation sequencing library qPCR kit and run on a Roche LightCycler 480 real-time PCR instrument. Sequencing of the flowcell was performed on the Illumina NovaSeq sequencer using NovaSeq XP V1.5 reagent kits, S4 flowcell, following a 2×151 indexed run recipe. Raw fastq file reads were filtered and trimmed using the JGI QC pipeline resulting in the filtered fastq file (*.filter-RNA.gz files). BBDuk (version 38.90) was used to remove contaminants (BBDuk, BBMap and BBMerge commands used for filtering are placed in file: 52510.2.364064.AAGAAGGC-AAGAAGGC.filter_cmd-MICROTRANS.sh), trim reads that contained adapter sequence and homopolymers of G’s of size 5 or more at the ends of the reads, right quality trim reads where quality drops below 6, remove reads containing 1 or more ‘N’ bases, remove reads with average quality score across the read less than 10, having minimum length <= 49bp or 33% of the full read length. Quality trimming was performed using the phred trimming method set at Q6. Finally, following trimming, reads under the length threshold were removed (minimum length 25 bases or 1/3 of the original read length - whichever is longer). The barcoded RNAseq libraries were loaded on one SP lane on an Illumina NovaSeq 6000 for cluster formation and sequencing. The libraries were sequenced from both ends of the fragments for 150bp from each end. The fastq read files were generated and demultiplexed with the bcl2fastq v2.20 Conversion Software (Illumina).

### 2.3 Plant growth conditions in soil for phenotypes and transcriptomics of nodules

To evaluate the effects of Hg on *M. truncatula* nodules and its effect on nodule number and plant biomass traits, we grew two *M. truncatula* genotypes (HM302, HM304) in sterilized turface as a soil substrate, which were inoculated with either of two *S. medicae* strains (AMp07, AMp08) that have different Hg-tolerance due to the presence of a mercury reductase operon in AMp08. The operon gives AMp08 a 10-fold increase in tolerance to Hg over AMp07 (Nonnoi et al., 2012). Plants were well-replicated (average of 20 plants per assay) and completely randomized. Prior to planting, seeds were treated in concentrated sulfuric acid for 3 minutes and washed 5 times with MiliQ water. Then seeds were surface sterilized using 50% bleach with 0.1% tween20 for 3 minutes and washed 8-10 times with sterilized water and left in sterile water for 2 hours at room temperature in dark. The seeds were placed on sterilized filter paper on petri dishes and kept overnight at 4°C in the dark. Later, the petri dishes were kept in a growth chamber at 21°C and 40% humidity with a photoperiod of 16/8 hours for a week and then seedlings were transferred to the sterilized Turface:vermiculite (2:1) soil (LESCO-turface all sport pro soil) in plant growth chamber. Plants were cultivated in a walk-in growth chamber with temperature set at 22°C and 16-h-light/8-h-dark photoperiod and watered every 5-7 days with sterilized ½ strength of B&D medium with low concentration of KNO_3_ (0.5 mM). Prior to inoculation KNO_3_ was not added while watering seedlings. Seedlings were inoculated with *S. medicae* AMp07 and AMp08 rhizobia strains 5-7 days after being transplanted to the sterilized turface soil. The inoculum was prepared by culturing rhizobia at 28°C in sterilized liquid TY medium until an OD600 of 0.8 was reached. The bacterial pellet was collected by centrifuging, washed and resuspended in 0.9% NaCl solution for the inoculation. At 21 days post inoculation (21dpi), plants were treated with ½ strength of B&D medium amended with 100μM HgCl_2_. Non-treated plants were watered with ½ strength of B&D medium supplemented with 0.5mM concentration of KNO_3_. Nodules were collected 7 days post treatment for phenotyping, which included quantification of fresh biomass, metal accumulation and nodule number, RNA extraction for RNA-seq, and XRF imaging. Nodules for RNA extractions were harvested, frozen in liquid nitrogen, and stored at −80°C.

### 2.4 Tissue sampling and isolation of RNA from nodules

To extract RNA from nodules, they were initially crushed in Qiagen TissueLyser II for 2 minutes. Thereafter, RNA was extracted using Spectrum Plant Total RNA kit (Sigma) by using the manufacturer’s protocol. Finally, the concentration and quality of RNA was quantified by using nanodrop (Thermo Scientific) and Bioanalyzer (Agilent) respectively.

### 2.5 Construction of strand specific RNAseq libraries from nodule samples

Construction of the RNAseq libraries and sequencing on the Illumina NovaSeq 6000 were performed at the Roy J. Carver Biotechnology Center at the University of Illinois at Urbana-Champaign. Purified DNased total RNAs were run on a Fragment Analyzer (Agilent) to evaluate RNA integrity. The total RNAs were converted into individually barcoded RNAseq libraries with the Universal Plus mRNA-Seq Library Preparation kit from Tecan, using custom probes against *M. truncatula* and *S. melliloti* rRNA designed by Tecan. Libraries were barcoded with Unique Dual Indexes (UDI’s) which have been developed to prevent index switching. The adaptor-ligated double-stranded cDNAs were amplified by PCR for 10 cycles. The final libraries were quantitated with Qubit (ThermoFisher) and the average cDNA fragment sizes were determined on a Fragment Analyzer. The libraries were diluted to 10nM and further quantitated by qPCR on a CFX Connect Real-Time qPCR system (Biorad) for accurate pooling of barcoded libraries and maximization of number of clusters in the flowcell.

### 2.5 Illumina read mapping, differential gene expression analysis and GO-enrichment

Illumina reads (150 bp) were trimmed using trimmomatic (Bolger et al., 2014). For leaf and root samples, Illumina reads were mapped to the *M. truncatula* (Mt5.0) reference genome (Pecrix et al., 2018) using STAR (Dobin et al., 2013). For nodule samples, the genome sequences of *M. truncatula* (Mt5.0) and *S. medicae* AMp07 or AMp08, and their gene annotations (.gff files) were concatenated to map plant and rhizobia reads to their respective genomes (“dual transcriptomics”). Indexed genomes of the concatenated genomes were made using STAR, and Illumina reads were mapped using STAR. Mapping and differential expression analysis of the AMp07 and AMp08 were reported in (Bhat et al., 2024). DESeq2 (Love et al., 2014) was used to quantify differentially expressed genes for leaf, root and nodule data using |log_2_ (Fold change) | ≥ 1 and false discovery rate (FDR) ≤ 0.05. We then used edgeR (Robinson et al., 2010) to quantify weighted trimmed mean of the log expression ratios (trimmed mean of M values (TMM)) (Robinson and Oshlack, 2010) as a normalized read count for measuring and visualizing relative gene expression values in Cd or Hg treated samples compared with control samples. We then incorporated functional annotation from Phytozome v13 for *M. truncaulta* v4.0. This annotation included KOG, KEGG, ENZYME, pathway, and InterPro annotations as well as the best BLAST hit to the *A. thaliana* genome. We merged the Mt5.0 with Mt4.0 gene annotations using the RNAseq dataset from Pecrix et al., (2018) in order to identify previously reported genes in literature predating Mtv5.0. We performed Gene-Ontology (GO) enrichment analysis on the differentially expressed genes using the PlantRegMap server (https://plantregmap.gao-lab.org/go.php) using species: *M. truncatula* and using default settings. Subsequently, dot plots were generated using gProfiler (https://anaconda.org/bioconda/gprofiler-official) within a custom python library using Python 3.9 Spyder. The dot plots were manually color-coded based on unique and shared responses observed under Cd and Hg stress.

### 2.6 Measurement of nodule number and biomass

The phenotypic data was collected 7 days post treatment (which equals 28 days post inoculation (dpi)). Plants were washed to remove surrounding substrate. The fresh biomass of shoot, root and nodule, and the nodule number were then measured to evaluate growth. Data analysis from different tissues (shoot, root, and nodules) was performed to compare *M. truncatula* genotypes (HM302 and HM304) inoculated with AMp07 and AMp08 rhizobia strains and to compare the differences between treated and non-treated plants. All experiments were performed in a randomized block design with at least 5 biological replications for each sample. Statistical significance for each data group was determined using student’s t-test with *p <*0.05.

### 2.7 Sample preparation and X-ray fluorescence (XRF) imaging

Nodules collected at 28 dpi were sectioned using a cryostat microtome (Leica CM1950). The nodules were mounted in O.C.T medium and longitudinally sectioned to 20µM thickness. The sections were mounted on a Ultralene thin film (3525) for XRF imaging. XRF images of nodules were acquired at the X-ray Fluorescence Microprobe beamline (4-BM) at the National Synchrotron Light Source II (NSLS-II). Briefly, this beamline uses Kirkpatrick-Baez (KB) mirrors to deliver focused X-rays (2-10mm spot) with tunable energy using a Si (111) double crystal monochromator. Samples were oriented 45° to the incident beam and the XRF detector (Canberra SXD 7-element SDD) was positioned 90° to the incident beam. Images were collected by continuously rastering the sample in the microbeam using a Newport stage with a 50mm pixel size and 50ms dwell time per pixel for course navigation maps, but a 5mm pixel size and 100ms dwell time per pixel for fine resolution maps. Data were processed using LARCH (Newville, 2013). Thin-film standard reference materials (NIST SRM 1832 & 1833) were measured as part of the data set to establish elemental sensitivities (counts per second per mg cm^−2^). The estimated ion concentrations were measured using a conversion of spectral images where Abundance (µg cm^−2^) = (CPS of the sample) x (Calibration ratio of element) / CPS of standard. In this equation, the term Abundance represents the concentration of the metal. The CPS of the sample refers to the average count per second measured, while the Calibration ratio refers to the standard for the specific element of interest. The “CPS of the standard” represents the value of the standards employed to normalize the concentration of the element of interest. Finally, the intensity was calculated by dividing the abundance by the thickness of the sample.

## 3 Results

### 3.1 The number of differentially expressed genes (DEGs) varies according to genotypes with different metal response phenotypes

The transcriptome for leaf and root tissues from the hydroponic experiment allowed us to compare DEGs among resilient and non-resilient classes of plant genotypes by screening a subset of *M. truncatula* genotypes from the HapMap collection. Four genotypes were selected for comparative transcriptomic analysis based on their Cd/Hg levels of leaf accumulation of either metal (Figure 1a, b): HM195 (low Cd accumulation), HM075 (high Cd accumulation), HM302 (low Hg accumulation) and HM304 (high Hg accumulation). Because all four genotypes exhibited low to intermediate RRG (i.e., low tolerance in roots) among the entire HapMap collection, we also included the genotype PI660407 which is a high Cd accumulator, shows high root tolerance to Cd, and had the highest root tolerance to Hg (García de la Torre et al., 2021, 2013).

**Fig. 1.**
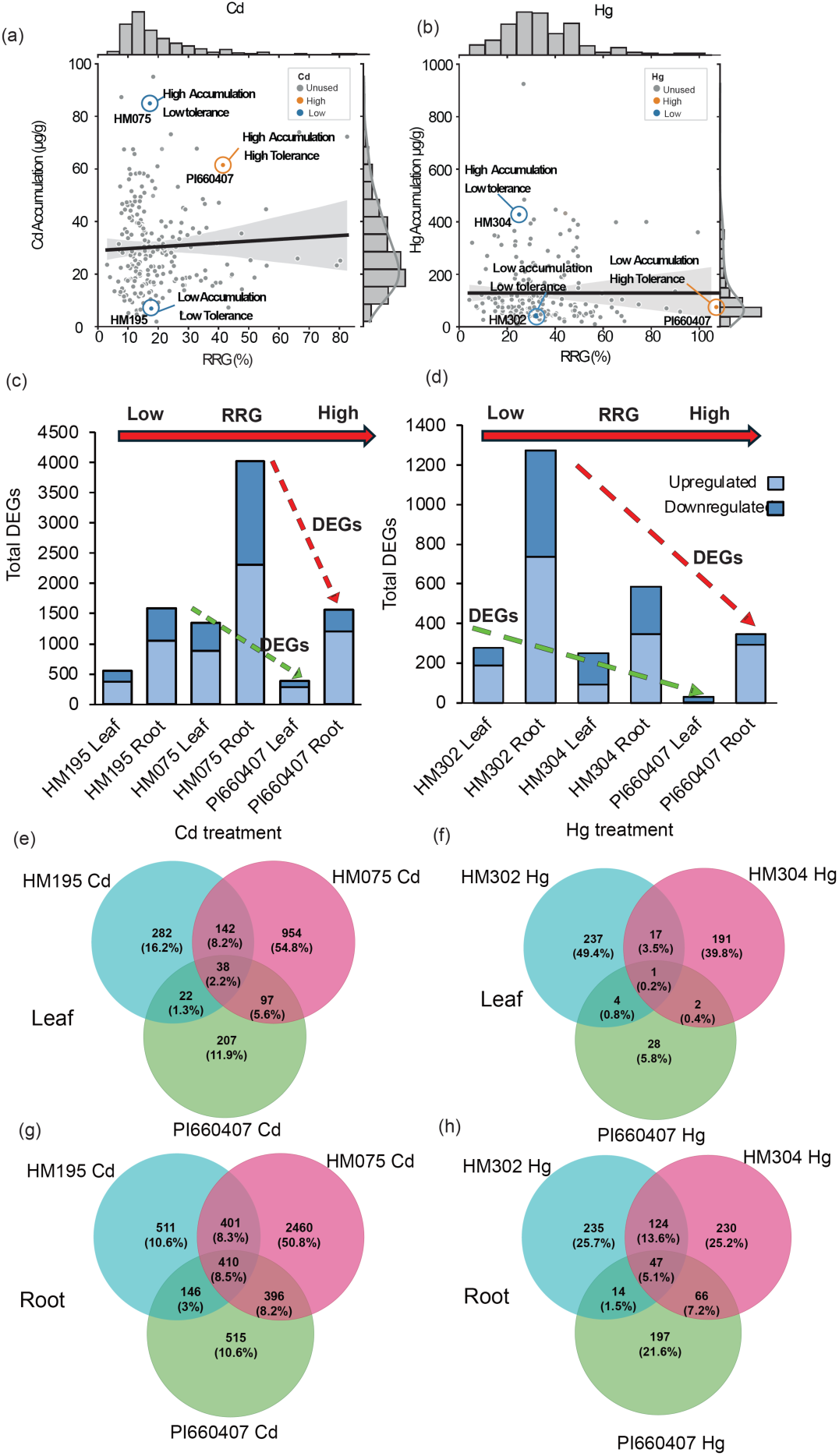
Scatterplot of leaf accumulation and relative root growth phenotypes of > 200 *Medicago truncatula* genotypes from the Medicago HapMap panel and USDA GRIN germplasm following treatment with 10μM CdCl_2_ (a) or 4μM HgCl_2_ (b) (Paape et al., 2022). The y-axis of (a) and (b) is accumulation of the metals in the leaf tissue, in units of µg/g of dry weight, and the x-axis is relative root growth (RRG) measured by comparing treated roots relative to untreated roots. The regression line shows the correlation between leaf accumulation and RRG for either metal treatment. The *M. truncatula* genotype names used in this study were labeled and colored orange or blue. All other genotypes measured but not studied here, are colored as gray points. The numbers of differentially expressed genes (DEGs) of the responding to Cd (c) and Hg (d) from plant genotypes in (a) and (b), where the y-axis is the counts of DEGs with light or dark blue shaded as up- or down-regulated, respectively. The x-axes of (c) and (d) are the plant genotype and tissue (leaf or root). The red arrow above the bar plots shows the genotype order in the x-axis from low to high RRG. The dashed red and green diagonal lines show the negative relationship between RRG and DEGs in roots and leaves, respectively. The Venn diagrams show the number of DEGs among the Cd-treated *M. truncatula* genotypes HM195, HM075, and PI660407 in leaf (e) and root (g) tissues that are unique or shared among the three genotypes, and the number of DEGs among the Hg-treated genotypes HM302, HM304, and PI660407 leaf (f) and root (h) tissues unique or shared among the three genotypes.

Plant genotypes exhibiting high tolerance to heavy metals may display a reduced number of DEGs as a result of their more precise responses to the stress. When considering RRG responses for either metal, this trend appeared to hold true as higher numbers of DEGs tended to occur in roots in the least tolerant genotypes, with the exception of HM195 which deviated from this trend for Cd-treated plants (Fig. 1c, d). There were also greater numbers of DEGs in roots compared with leaves for all genotypes, which was not surprising as roots make the first contact with the stress condition. The two high Cd accumulation genotypes (HM075 and PI660407) shared only a small portion of DEGs with 3.5% and 14.4% common genes in leaf and root tissue, respectively (Figure 1 e, f). For Cd treated plants, the Cd tolerance/accumulation phenotype was only partially predictive of the number of DEGs, but overall the most resilient genotype for tolerance/accumulation did have the fewest DEGs.

For Hg treated plants, the trend of fewer DEGs in more resilient genotypes was clearer. The number of DEGs followed the order for both root and leaf tissues: low Hg accumulator (HM302) > high Hg accumulator (HM304) > Hg-tolerant (PI660407) (Figure 1d, Table S2). The pattern suggests the more resilient genotypes (PI660407 and HM304) might have more efficient Hg detoxification mechanisms which results in reduced stress responses and therefore fewer DEGs. Interestingly, relatively small percentages of genes were shared (0.6% in leaf and 12.3% in root tissue) between the two most resilient genotypes (HM304 and PI660407) and the majority of DEGs were unique to either of these genotypes. This suggests they have evolved different mechanisms that contribute to their resilience to mercury stress. Conversely, there were substantially more shared DEGs between the low and high accumulator genotypes (HM302 and HM304) respectively (10.4% in leaf and 13.8% in root tissue) (Figure 1 g, h), despite substantial differences between these two genotypes in their response to Hg in root tissues.

### 3.2 Enrichment of gene families reveals both shared and genotype-specific responses to heavy metals

To determine whether plant genotypes responded to heavy metal stress using shared or genotype-specific DEGs, we used GO-enrichment to identify patterns of metal-specific and genotype-specific molecular functions (MF) and biological processes (BP) using the significant DEGs that responded to Cd and Hg stress (Supplementary Data File 2). First, we looked at GO-terms that were shared among the two metal treatments and found clustering of shared responses. These included response to oxidative stress, cellular oxidant detoxication, peroxidase activity, glutathione transferase activity, response to metal ion and UDP-glycosyl transferase (UGTs) activity which were enriched in all genotypes for both Cd and Hg stress, which suggests conserved molecular responses for Cd and Hg stress among diverse genotypes (Figure 2). The shared GO-terms for Cd and Hg stress show expected ontologies that reflect cellular and molecular responses to manage heavy metal toxicity in plant cells and tissues (Moustakas, 2023; Paape et al., 2022; Sharma and Lenaghan, 2022; Singh et al., 2016).

**Fig. 2.**
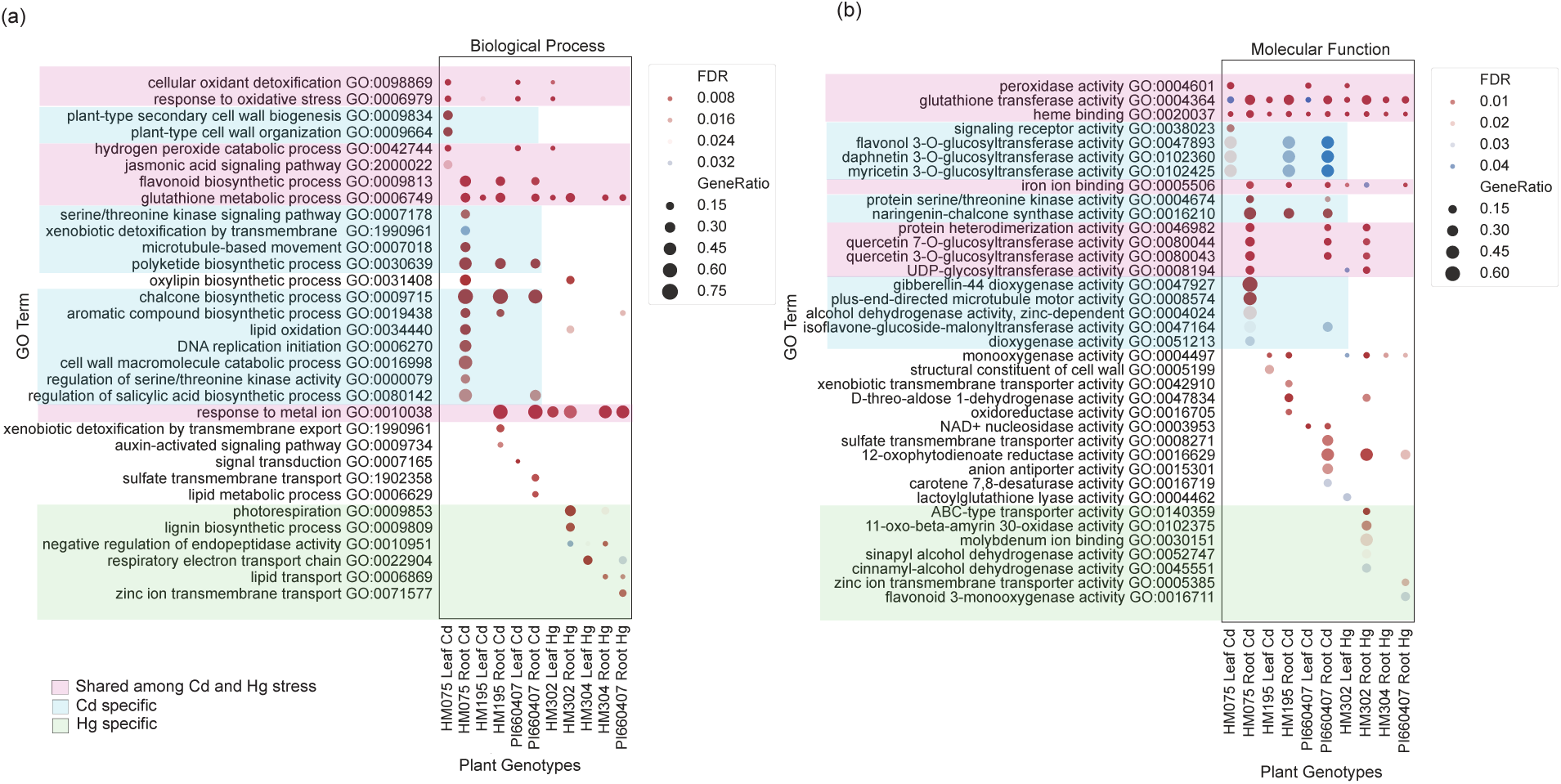
Gene ontology (GO) enrichment analysis of DEGs in leaf and root tissues of *M. truncatula* genotypes showing Cd treated genotypes HM075 (high accumulation) and HM195 (low accumulation) and the Hg treated genotypes HM302 (low accumulation) and HM304 (high accumulation). The dot-plot shows enriched GO-terms for DEGs using biological processes (a) and molecular functions (b) where the y-axis is the name of GO-terms along with their GO IDs. DEGs used for GO-enrichment were defined as log_2_FC ≥ 1 and adjusted p-value < 0.05. The x-axes of (a) and (b) are the plant genotypes, metal treatment and tissue (leaf or root). The color of the circle indicates the false discovery rate (FDR) adjusted p-value of over-enriched GO-terms. The gene ratio is the percentage of total DEGs in the given GO-term and circle size corresponds to relative abundance of genes in that GO term. We grouped sets of GO-terms with light-pink shaded areas corresponding to shared GO-terms between Cd and Hg treated genotypes, light-blue shaded for GO-terms associated with only Cd treated genotypes and light green shaded for GO-terms associated with only Hg treated genotypes.

Next, we examined expression patterns of genes within key GO-terms relevant to Cd stress (Figure S1), which included UGTs, oxidative stress response, glutathione-S-transferase activity and response to metal ion GO-terms (Figure S2, Supplementary Data File 2). UGTs are enzymes responsible for facilitating the chemical modification of diverse organic and inorganic compounds, and have been shown to respond to various abiotic stress conditions as well as interacting with glutathione to mediate oxidative stress (Blanco-Herrera et al., 2015; Caygill and Dolan, 2023; Meech et al., 2019). For UGTs, we observed distinct patterns of regulation characterized by clustering of downregulated and upregulated genes (Figure S2a), with the strongest genotype specific response shown in the high accumulating genotype. Oxidative stress is a ubiquitous metal ion stress response in plants which is characterized by heightened production of ROS and/or compromised antioxidative defense mechanisms (Cuypers et al., 2023). Genes enriched in oxidative stress response were extracted from GO-terms related to cellular antioxidant detoxification, response to oxidative stress, and peroxidase activity (Figure S2b). These genes had functional annotations that included superoxide dismutase (SOD), catalase (CAT), and peroxidases (POD) with a large clustering of downregulated genes, and a smaller cluster of upregulated genes in the lower highlighted section of the heatmap (Figure S2b).

Glutathione S-transferases are a super-family of enzymes that play important roles in oxidative stress responses induced by heavy metals by forming conjugates with GSH, which then can transport excess metals into vacuoles (Yadav, 2010). The glutathione S-transferase activity GO-term was enriched for all genotypes, and S-transferase genes showed up-regulation in nearly all genotypes (Figure S2c). The direct role of glutathione conjugates and glutathione S-transferases in heavy metal stress responses is supported by the nearly universal upregulation in response to Cd stress in both leaf and root tissues. Notably, genotype-specific responses were not very obvious based on the glutathione S-transferase gene expression, and it is likely that co-expressed glutathione S-transferase genes work collaboratively to detoxify leaf and root tissues based on their expression patterns. Furthermore, genes associated with the response to metal ion GO-term were identified as glutathione S-transferase family genes and mostly showed upregulation in roots in all genotypes (Figure S2d).

We used a similar approach to visualize Hg-dependent genotypic responses of genes associated with ABC-transporter activity, glutathione-S-transferase activity, UDP-glycosyltransferase activity, oxidative stress and response to metal ion GO-terms (Figure S3). Notably, the genotype characterized by low Hg accumulation (HM302) demonstrated an enrichment in GO-terms related to response to oxidative stress, peroxidase activity, cellular antioxidant detoxification, ABC-transporter activity, UDP-glycosyltransferase activity, and photorespiration (Figure S3 a, b). This suggests a multifaceted response to Hg stress in the low Hg accumulator, indicative of a broader spectrum of DEGs associated with Hg stress. In contrast, the high Hg accumulating genotype (HM304) showed enrichment in GO-terms specifically linked to detoxification pathways, such as heme binding and glutathione transferase activity. This pattern suggests that high accumulators may employ an alternative and more selective mechanism for Hg stress response. Unlike the Cd response gene expression patterns, the genes in the ABC-transporter activity, glutathione-S-transferase activity, UDP-glycosyltransferase activity, oxidative stress and response to metal ion GO-terms showed much stronger genotypic response, particularly for the low accumulator (HM302) genotype (Figure S4).

### 3.4 Several genes identified using genome wide association studies showed differential expression

We found that many of the candidate genes identified using GWAS (Paape et al., 2022) showed genotype specific responses, as expected if the associations were driven by allelic differences. Among these genes included are MtABCC2, MtABCC3, MtNRAMP6, MtHMA2, MtCAX3, ankyrin repeat protein, Fe-OG, H^2+^ ATPAse2 (Figure S5). Most of these genes belong to well described gene families that involve micronutrient uptake and transport, and toxic ion compartmentalization (Brunetti et al., 2015; Cheng et al., 2005; Do et al., 2021; Park et al., 2012). While the findings from GWAS also showed highly polygenic associations to the heavy metals, the transcriptomics datasets support that Cd/Hg tolerance and accumulation are regulated by polygenic factors which depend on transport, compartmentalization, and oxidative stress responses, but we could more comprehensively identify gene family responses as described in the previous sections.

### 3.5 Mercury tolerant rhizobia affects heavy metal stress responses in host plants

Heavy metal resistant symbiotic rhizobia strains have been isolated from legumes host plants that grow in contaminated mines (Fagorzi et al., 2018; Lu et al., 2017). We inoculated two plant genotypes that showed phenotypic differences in Hg accumulation and tolerance, and in their transcriptomes in the hydroponic experiment (HM302 and HM304 (Figure 1b, d), with two *S. medicae* strains with low Hg tolerant (“LT”, AMp07) or high Hg tolerant (“HT”, AMp08), then treated the plants with Hg stress. The AMp08 strain possesses a mercury reductase (Mer) operon (Bhat et al., 2024), which allows it to grow at a minimum inhibitory concentration (MIC) of 250µM of HgCl_2_ (Bhat et al., 2024; Nonnoi et al., 2012), which is a 10-fold difference in Hg tolerance compared with the LT AMp07 strain which does not possess a Mer operon.

Among the different host plant-rhizobia strain combinations, HM304-AMp08 showed the highest shoot biomass in control conditions (Fig. 3a). However, root biomass for this same host-strain combination was significantly lower than when strain AMp07 was used (Fig. 3b), suggesting resource allocation to shoots. Nodule biomass was also highest for HM304-AMp08 but not significantly (Fig. 3c, Fig. S7), while nodule number and nodule number/root biomass significantly higher for HM304-AMp08 (Fig. 3d-e). These results suggest strain-host compatibility for plant growth promotion in the absence of Hg stress. For all phenotypes except nodule number/root biomass, Hg treatment significantly reduced the traits in each host-strain combination. We also found that nodule number after Hg treatment in HM304-AMp08 was significantly higher than HM304-AMp07 (Fig. S7). The Hg-tolerant combination HM304-AMp08 was the least affected by the Hg stress in shoot biomass.

**Fig. 3.**
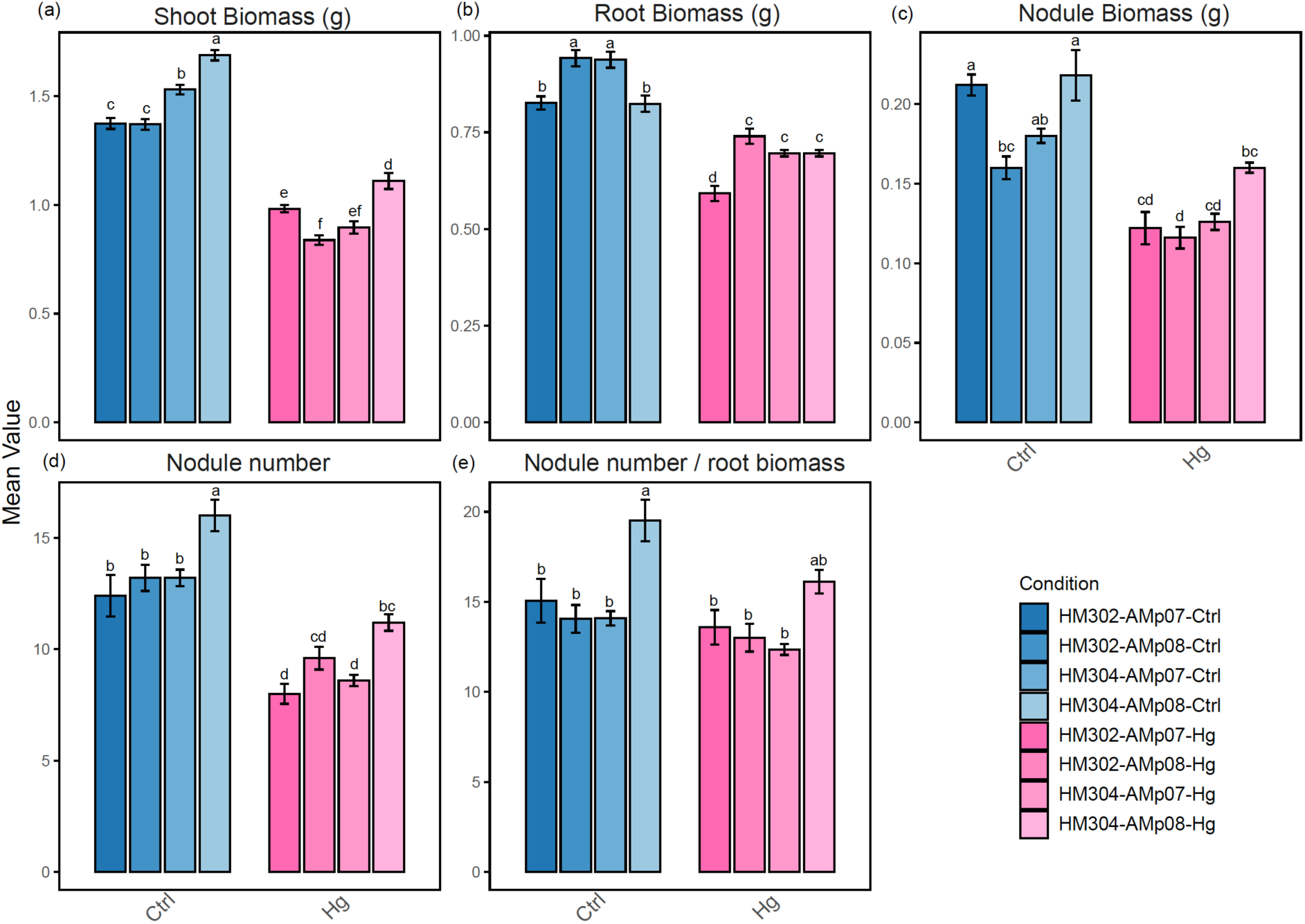
Biomass measures for shoot (a), root (b), and (c) nodule in control (blue hues) and Hg treated (red hues) *M. truncatula* plants (y-axes are in grams). Five plants were tested in each condition, and the experiment repeated three times. Nodule number (d) is the total number of nodules among five independent replicates (y-axis). The number of nodules indicates total nodules pooled from five plants. Nodule number per biomass (e) is total nodules normalized by root fresh weight (y-axis). The legend shows the different host plants (HM302 and HM304) and the respective S. medicae strain used as inoculants (AMp07 “LT”, and AMp08 “HT”). Error bars indicate standard error of the mean (SEM). Letters display Tukey’s Honestly Significant Difference (HSD) test for multiple comparisons at α = 0.05. When common letters are shown above any bar, they are not significantly different.

Alteration of micronutrient distributions is known to be affected by heavy metals in legumes (Pandey et al., 2022), most likely as the result of increased oxidative stress that disrupts ion homeostasis. Given the crucial role of iron (Fe) and zinc (Zn) in nodule formation (Brear et al., 2013; Kryvoruchko et al., 2018; León-Mediavilla et al., 2018), we used X-ray Fluorescence (XRF) imaging to compare the distribution of Fe and Zn in the nodules harboring either LT or HT *S. medicae* strains. In control conditions, XRF imaging revealed no visual differences in the distribution of Fe and Zn among the genotype and host-strain combinations, but both Fe and Zn levels were visually reduced following Hg treatment in both host genotype and rhizobia strain combinations (Figure 4 a, c). The visual differences were confirmed by quantification of elemental abundance which showed a significant reduction in Fe abundance in both plant genotypes after Hg treatment (Figure 4 b, d). Higher Fe abundance was found when the HT strain was used compared to plants inoculated with the LT strain after Hg treatment (Figure 4 b, d). No significant difference was observed in Zn abundance in Hg treated nodules compared with controls. Our results suggest that the Hg-tolerant rhizobia strain may directly or indirectly reduce the impact of Hg stress in the host plants which thereby reduces the impact on Fe.

**Fig. 4.**
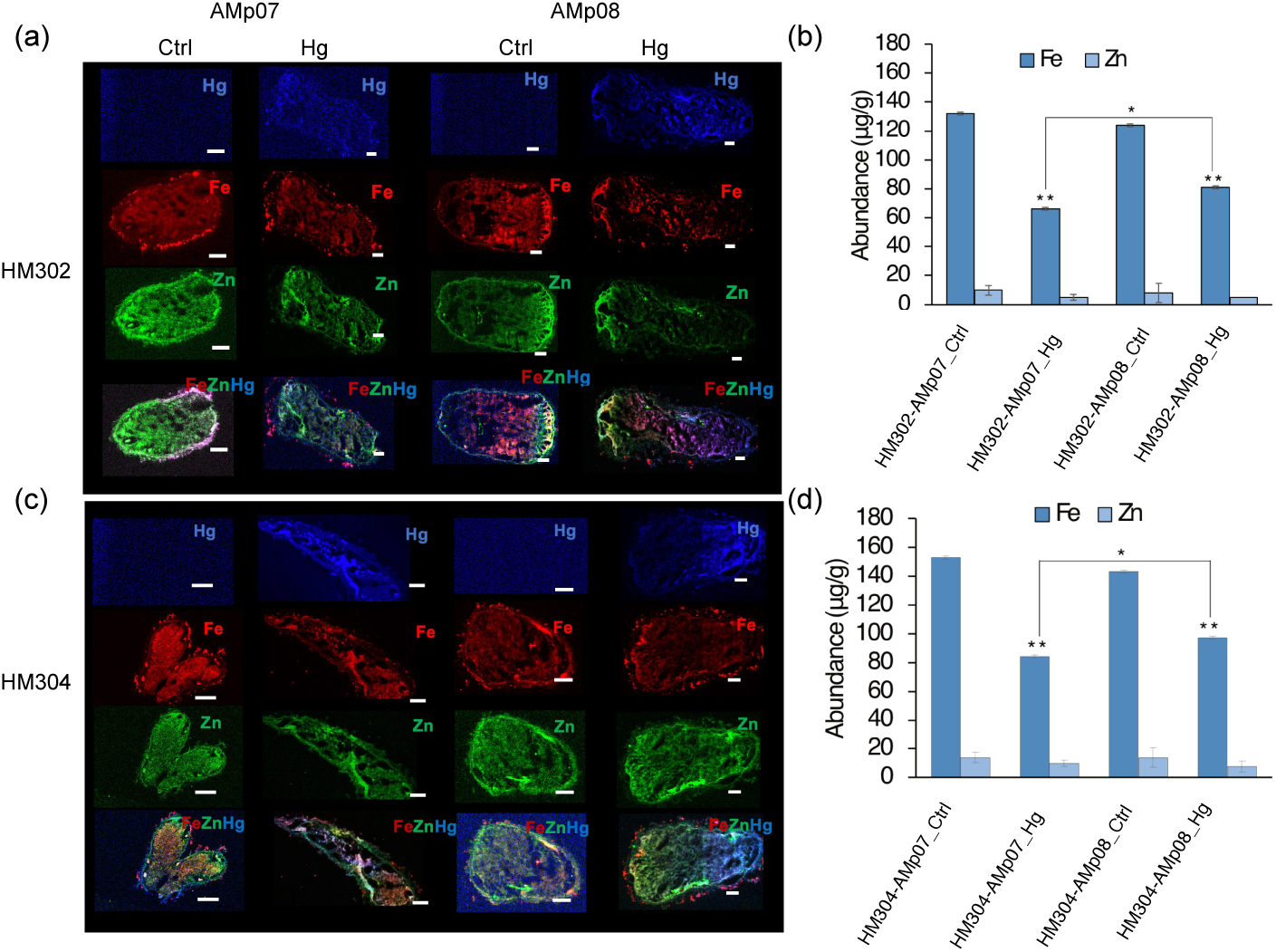
(a) Spatial distribution of mercury (Hg), iron (Fe), and zinc (Zn) within nodule cross sections (20μm) of HM302 genotype inoculated with the *S. medicae* strains AMp07 and AMp08 of non-Hg treated (Ctrl) and 100μM Hg treatment (Hg). Images of ion distributions were measured using X-ray fluorescence (XRF). (b) Metal abundance of Fe and Zn within nodule cross sections of the plant genotype HM302. Quantification of elemental abundance of Fe and Zn were calculated by using a conversion of spectral images to Abundance (μg/g). (c) Spatial distribution of Hg, Fe, and Zn within nodule cross sections of HM304 for control and Hg treated plants, inoculated AMp07 and AMp08. (d) Metal abundance of Fe and Zn within nodule cross sections of the plant genotype HM304. For abundance comparisons in b and d, statistical comparisons of Fe or Zn were made between Ctrl and Hg. The black bar represents a statistical comparison of Fe between AMp07 and AMp08 inoculated HM302 and HM304 host plants after Hg treatment in (b) and (d). The bar graphs show the mean ± SD of 3 biological replicates, and significance was calculated using Student’s t-test (**p < 0.01, *<0.05). The scale bar is 1mm in Figures (a) and (c).

### 3.6 Transcriptomic response in host plants during symbiosis is determined by rhizobia tolerance to Hg

The transcriptomic response in the nodules showed a clear reduction in the number of DEGs in both host plant genotypes in symbiosis with the HT strain, as nodules with the LT strain had 2-3 times higher numbers of DEGs compared to the HT strain (Figure 5a, Table S2). This finding indicates that when plants form nodules with the more Hg-tolerant *S. medicae* strain, the strain plays a role in mediating or dampening the stress response, consistent with the phenotypic comparisons reported above. The volcano plots visually represent the distributions of up- and down-regulated DEGs (Figure 5 b-e), where labeled genes indicate those that have been significantly affected by Hg stress among different host genotype and rhizobia combinations (Table S3-S6). In the HM302-AMp07 combination (low accumulating plant genotype-low tolerant rhizobia), the downregulation of Zn family transporter genes suggests prohibiting the uptake of Hg ions via essential metal ion transporters (Table S3). This indicates a plant strategy to limit the influx of additional Hg and detoxification processes mediated by upregulated ABC-transporters. Additionally, the upregulation of UGTs suggests their involvement in the conjugation of Hg ions with compounds such as phytochelatins, facilitating their sequestration or detoxification. In the HM302-AMp08 (low accumulating-high tolerant) combination, the most significantly upregulated genes are related to glutathione and oxidative stress, which suggests an antioxidant defense system in response to Hg stress. This response likely helps mitigate the oxidative damage caused by Hg exposure, supporting the survival and growth of the low Hg-accumulating plant genotype when inoculated with the Hg-tolerant strain. Furthermore, the upregulation of UGTs in this combination indicates a concerted effort to conjugate and detoxify metal ions (Table S4).

**Fig. 5.**
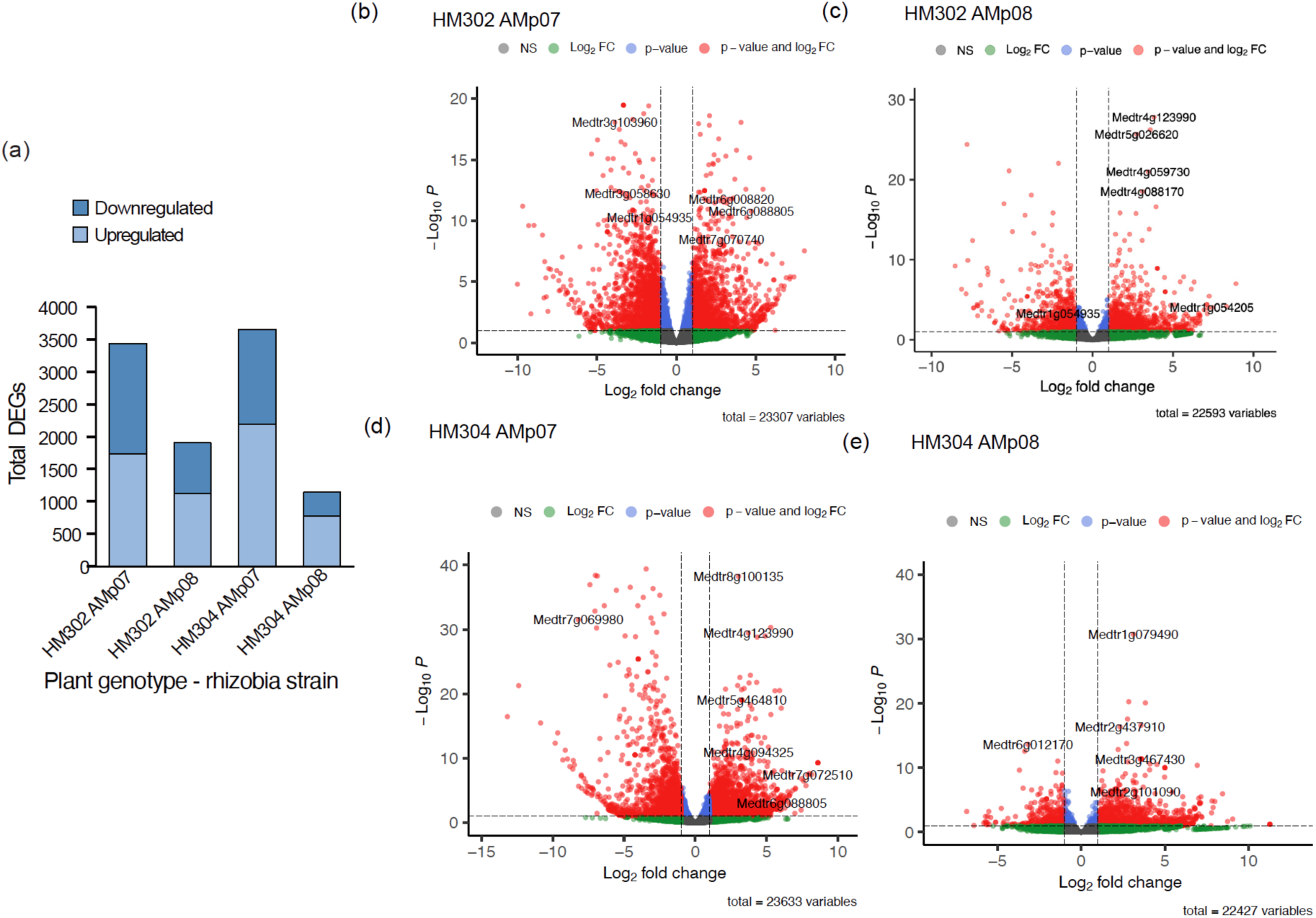
The number of DEGs of the HM302 and HM304 genotypes inoculated with AMp07 and AMp08 in nodules after Hg treatment (a) where the y-axis is the total number of DEGs with light or dark blue shaded as up-or down-regulated, respectively. The x-axis is the plant genotype and rhizobia strain combination. Volcano plots of DEGs from (b) HM302-AMp07 (c) HM302-AMp08 (d) HM304-AMp07 (e) HM304-AMp08 plant genotype and rhizobia strain combination. The x-axis is the fold change and on the y-axis the p-value. Gray represents not-significant change in expression whereas red represents significant DEGs with log_2_FC ≥ 1 and p-value < 0.05, blue represents p-values and green represents the log_2_FC. The gene IDs correspond to significant genes belonging to categories such as metal transporters and response to oxidative stress, which are predominant in our GO enrichments (Table S3-S6).

In the HM304-AMp07 combination (high accumulating-low tolerant), we found strong upregulation of genes associated with oxidative stress, UGTs, and vacuolar transport. Because this combination is with a high Hg-accumulating plant genotype inoculated with a LT strain, these genes indicate both detoxification mechanisms mediated by UGTs and vacuolar transport, as well as antioxidant defense systems to mitigate Hg-induced oxidative damage (Table S5). Finally, in the HM304-HT (high accumulating-high tolerant) combination, the upregulation of ABC transporters, glutathione associated genes, UGTs, and oxidative stress represents a comprehensive response to Hg stress. This response likely facilitates efficient Hg detoxification and sequestration while minimizing oxidative damage through enhanced antioxidant defenses (Table S6).

### 3.7 GO-enrichment shows plant genotype by strain specific molecular functions and biological processes

Using GO-enrichment analysis of genotype and strain combinations we found clear clustering of GO-terms associated with groups of biological processes. These included four general categories: a) cell division, cell cycle and transmembrane transport, b) abiotic stress and hormone signaling, c) response to oxidative stress, and d) ABC-transporter, glutathione, peroxidase (Figure 6). While these groupings are fairly broad and not entirely mutually exclusive, they showed that each of the four plant genotype-rhizobia strain combinations comprised different sets of gene ontologies, which indicated that the different symbiotic combinations responded to Hg stress very differently at the transcriptome level. The category of cell division, cell cycle and transmembrane transport (pink shading in Figure 6) includes essential processes for basic cellular function including sulfate and sugar transmembrane transport, anion transporters, DNA replication, carbohydrate metabolism, response to water deprivation, microtubule binding and protein kinase activity. These GO-terms suggest basic housekeeping mechanisms were disrupted when the LT strain was used in both of the plant genotypes. Interestingly, the response to oxidative stress group of GO-terms (green shading in Figure 6) was limited to the nodules of the high Hg accumulating genotype with either rhizobia strain. This was somewhat surprising since oxidative stress is a ubiquitous response to heavy metal stress in plants, and the low Hg accumulating genotype, HM302 showed no enrichment for these GO-terms. The abiotic stress group of GO-terms (light blue in Figure 6), which also included hormone signaling, was almost exclusively limited to the low Hg accumulating plant genotype (HM302) in symbiosis with the HT strain By contrast, the high Hg accumulating plant genotype (HM304) in symbiosis with the HT strain showed GO-terms that are commonly associated with heavy metal transport, heavy metal stress and detoxification including ABC-transporter activity, glutathione transferase activity, UDP-glycosyltransferase activity, and iron and heme binding. The pattern of gene expression for ABC-transporter activity, glutathione transferase activity, and UDP-glycosyltransferase activity genes showed the strongest up and down-regulation in the HM304-HT combination which reflects well the GO-enrichment patterns for this plant-strain combination (Figure S8).

**Fig. 6.**
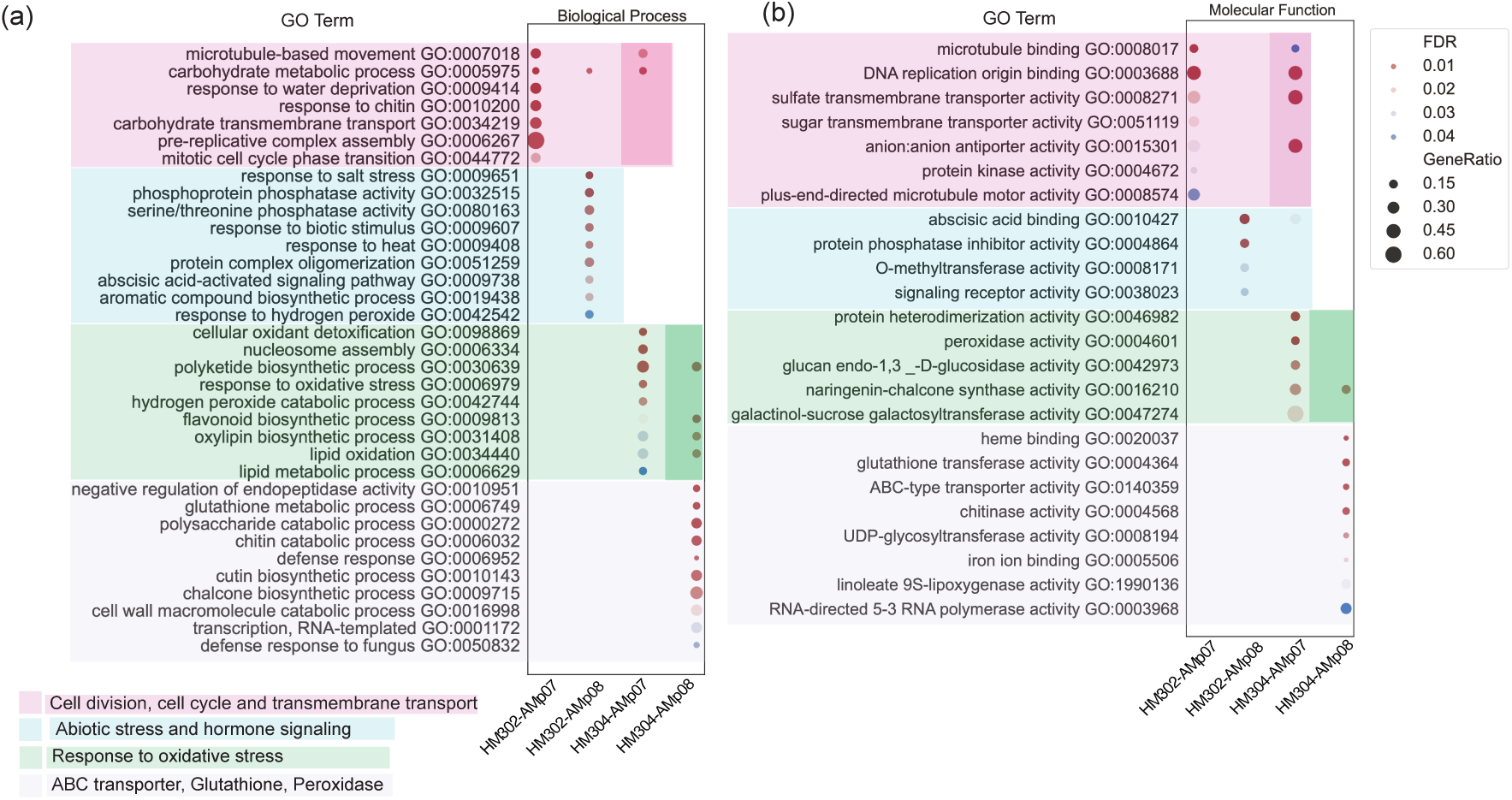
GO-enrichment analysis of DEGs from Hg-treated nodules of *M. truncatula* genotypes HM302 and HM304 inoculated with *S. medicae* strains AMp07 and AMp08. Dot-plots show enriched GO terms for biological processes (a) molecular functions (b), where the y-axis is the name of GO-terms and GO-term ID. Light pink shaded areas indicate cell division, cell cycle and transmembrane transport related GO-terms, the light blue shaded areas indicate abiotic stress and hormone signaling related GO-terms, light green shaded areas indicate response to oxidative stress related GO-terms, and light purple shaded areas indicate ABC-transporter, glutathione, and peroxidase related GO-terms. The shading of these GO-terms highlights different hostplant and rhizobia strain combinations have different biological processes and molecular functions. The dark pink and dark green shows that some of the highlighted GO-terms are shared by two hostplant and rhizobia strain combinations. The x-axes of (a) and (b) are the plant genotypes and rhizobia combinations. The dotplot enrichment legend indicates the color of the circle is the adj. p-value (FDR) of the over-enriched GO-terms, the gene ratio is the percentage of total DEGs in the given GO term and circle size corresponds to relative abundance of genes in that GO term.

### 3.8 Fewer plant symbiosis genes responded to Hg stress when inoculated with Hg tolerant rhizobia

Previous research has characterized a large number of plant genes that play crucial roles in one or more stages of symbiosis and nodulation (Hohnjec et al., 2009; Jardinaud et al., 2016; Lee et al., 2024; Roy et al., 2020; Zhang et al., 2022). We analyzed the differential expression of plant genes that have been previously implicated in legume-rhizobium symbiosis and observed fewer number of DEGs in plant genotypes inoculated with the HT (AMp08) strain compared to the low tolerant LT (Amp07) strain (Figure 7a), suggesting less impact of Hg stress on symbiosis genes in the host plant when inoculated with the Hg tolerant rhizobia strain possessing a Mer operon (Bhat et al., 2024). The least number of total DEGs was observed in the HM304-AMp08 pair, followed by the HM302-AMp08 pair which is consistent with the genome wide pattern (Figure 5a). Moreover, the HM304-AMp08 pair had nearly equal up and downregulated genes, but the other three host plant-rhizobia combinations had 2-3 times more upregulated DEGs compared to those that were downregulated (Figure 7a, Supplementary Data File 3). Among the symbiosis gene subset, less than 20% of the significant genes were shared between the two host plant genotypes when the LT strain was used, and only 3 DEGs were shared between the two host plant genotypes when the high tolerant rhizobia strain was used (Figure 7b). Despite the small number of DEGs shared across the four different plant-rhizobia combinations due to statistical thresholds of counting DEGs, the HM304-AMp08 combination of high accumulator plant, high Hg tolerant rhizobia, respectively, clearly stands out as the most dissimilar with the other three combinations sharing large numbers of symbiosis genes with common expression patterns (Figure 7c).

**Fig. 7.**
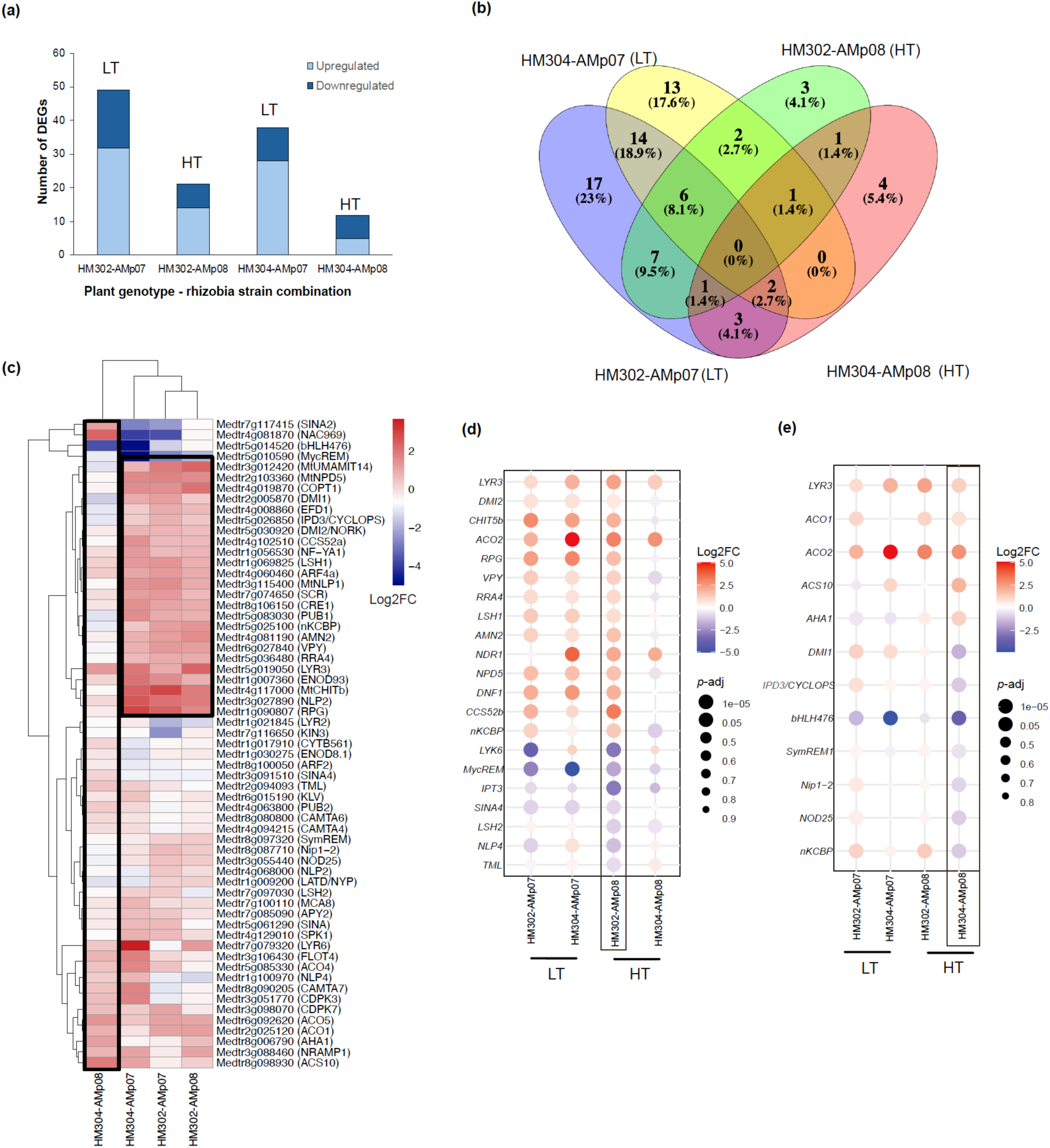
Differential expression of known symbiosis related genes in *M. truncatula*. (a) Number of symbiosis genes differentially expressed under Hg stress in nodules in all four combinations of host plant genotypes and rhizobia strains. Bars were colored red and blue upregulated or downregulated genes, respectively. (b) Venn diagram showing the shared DEGs among four different plant genotype-rhizobia strain combinations. (c) Heatmap showing the expression of symbiosis genes in Hg treated vs. control conditions where red is upregulated, and blue is down regulated. The log2FC values were clustered on the x and y-axes, shown by the dendrograms. The black around the HM304-AMp08 combination highlights the different expression patterns compared with the other three combinations. The upper right-side black box highlights a large set of upregulated symbiosis genes in the other three combinations. (d) Symbiosis genes differentially expressed after Hg stress in nodules in the HM302-AMp08 combination, which may or may not be differentially regulated in the three other host plant genotype and rhizobia strain combination. (e) Symbiosis genes differentially expressed after Hg stress in nodules in the HM304-AMp08 combination, which may or may not be differentially regulated in the three other host plant genotype and rhizobia strain combinations. Only genes with expression levels corresponding to p-adj <0.05 were considered differentially regulated in the HM302-AMp08 (d) or HM304-AMp08 combination (e). The red-blue scale bar to the right of (c), (d) and (e) is log2FC where red corresponds to upregulated, and blue corresponds to downregulated. Dot sizes in (d) and (e) are shown according to their adjusted p-value (p-adj) where larger dots are smaller p-adj values. HT and LT refer to high tolerant (AMp08) or low tolerant (AMp07) strains, respectively.

When the LT strain was the symbiont in the HM302 and/or HM304 genotypes, the largest number of symbiosis genes were upregulated including multiple *LYKs* and *LYRs*, and early signaling genes such as the calcium-dependent calcium channel-encoding *DOES NOT MAKE INFECTION 1 (DMI1)* and the receptor-like kinase-encoding *DMI2.* Additionally, there was upregulation observed in *MtCHITb*, a gene encoding a hydrolase involved in degrading nodulation factors (NFs). Furthermore, early signaling genes such as *M. truncatula CALCIUM ATPase FAMILY 8 (MCA8),* and *PLANT U-BOX PROTEIN 1 (PUB1*), infection-associated *EPS RECEPTOR 3 (EPR3/LYK10), RHIZOBIUM-INDUCED POLAR GROWTH (RPG), FLOTILLIN 4 (FLOT4*), and *VAPYRIN (VPY),* transcription factor-encoding *NUCLEAR FACTOR-YA1 (NF-YA1)* involved in both infection and nodule organogenesis, cell division-associated *SCARECROW (SCR)*, nodule organ identity-related *LIGHT-SENSITIVE SHORT HYPOCOTYL 1 (LSH1*), and nodule differentiation-associated CELL CYCLE SWITCH 52 *(MtCCS52), DEFECTIVE IN NITROGEN FIXATION 1 (DNF1)* and *M. truncatula NATURAL RESISTANCE-ASSOCIATED MACROPHAGE PROTEIN 1 (MtNRAMP1*). Genes such as *EPR3, RPG,* and *VPY* have been previously observed to be upregulated under salt stress (Chakraborty et al., 2021), suggesting that these symbiotic genes are particularly sensitive to environmental fluctuations. The cytokinin receptor-encoding *CRE1,* the direct target of cytokinin, *CYTOKININ RESPONSE REGULATOR (RRA4),* and multiple ethylene synthetic genes, *ACC synthase (ACS*) and *ACC oxidase (ACO*) were upregulated among several other genes, implicating these hormones in responding to Hg stress in the nodules (Figure 7c, Supplementary Data File 3). The large number of differentially expressed symbiosis genes in response to Hg stress with the LT strain are likely compensating for greater impacts of the stress on nodulation compared with plants inoculated with the HT strain (see Figure 3b).

Focusing on the symbiosis DEGs in the host plants inoculated with the HT strain (AMp08), we found a strong plant genotype effect where the HM302 and HM304 symbiosis gene sets show considerable differences despite their inoculation with the HT strain (Figure 7d, e). For the significant DEGs in the low Hg accumulating HM302 genotype, the majority were upregulated, and their expression was more similar to expression in host plants that were inoculated with AMp07 where most of the same genes also showed upregulation in these conditions (Figure 7d). In the set of DEGs from the HM302-AMp08 pair, a few of these genes, such as *LYR3, DMI2*, *CHIT5b, RPG, VPY, LSH1, CCS52b,* and *DNF1* were still observed. Also upregulated in this pair were ethylene synthetic genes *ACO2*, and the symbiotic *ABC-*transporter-encoding *AMN2* (Figure 7d, Supplementary Data File 3). The Hg tolerant HM304-AMp08 pair had the least number of DEGs, and other than *ACO2* and *LYR3,* none of the symbiotic DEGs observed in the HM302-AMp08 pair were differentially regulated in this pair (Figure 7d, Supplementary Data File 3). On the contrary, *ACC synthase 10 (ACS10),* which is suggested to be a mediator of nitrate-inhibition of nodulation, and the plasma membrane proton pump-encoding *MtAHA1* that energizes nutrient uptake during mycorrhizal symbioses were uniquely upregulated in this pair (Figure 7d, Supplementary Data File 3).

The Hg tolerant HM304-AMp08 pair also showed several uniquely downregulated symbiotic genes, which were higher in number than the upregulated genes in this category. The set consisted of the DEGs that were either not observed in any other pair or were upregulated in one or more of the other pairs. This set of uniquely downregulated genes includes *DMI1* and *INTERACTING PROTEIN OF DMI3 (IPD3)*. IPD3 plays a critical role during the initiation of rhizobium-legume symbiosis by activating several downstream transcription factors. In addition, the infection-associated remorin-encoding *SymREM1,* the multifunctional channel-encoding *NOD26-LIKE INTRINSIC PROTEIN (nip1-2),* differentiation-associated *NODULIN 25 (NOD25),* and *NODULATION-SPECIFIC KINESIN-LIKE CALMODULIN-BINDING PROTEIN (nKCBP)* were also uniquely downregulated in this Hg tolerant HM304-AMp08 pair, revealing a general trend of downregulation, or minimal perturbation of genes playing critical roles at various stages of nodule development (Figure 7c, Supplementary Data File 3). Notably, *DMI1, IPD3, Nip1-2,*and *nKCBP* were significantly upregulated in the HM302-Amp07 and / or in the HM304-Amp07 pairs, where the host was low Hg accumulator (Supplementary Data File 3). These results suggest that the Hg tolerance in both the host and rhizobia combination contribute towards the lower expression of symbiotic genes in the nodules under Hg stress.

## 4 Discussion

### 4.1 Heavy metal stress in *M. truncatula* genotypes in leaves and roots

Genetic diversity is key to identifying adaptive mechanisms to stress in plants. Resilience to Cd and Hg stress in plants is genotype specific in non-hyperaccumulating plants and linked to inducible genetic responses (Alvarez-Rivera et al., 2022; Chao et al., 2012; García de la Torre et al., 2021, 2013) as our phenotype data and plant transcriptome analyses indicated. First, in the hydroponic experiment, the majority of DEGs were unique for each genotype for both metal treatments, with only small percentages of shared DEGs between genotypes in either Cd or Hg treated genotypes. This suggests that even if a gene of large effect that could transport both Cd and/or Hg into vacuoles such as ABC-proteins and ATPases as in *A. thaliana* (Chao et al., 2012; Kang et al., 2011; Park et al., 2012), there will likely be other polygenic differences among genotypes in their transcriptomic responses. Second, we did not observe a strong correlation between the number of DEGs and metal accumulation in Cd accumulators, while all genotypes of high Hg accumulators showed lower numbers of DEGs when compared to their low accumulator counterparts, a pattern more consistent with the relationship between resilience and stress responses in the transcriptome. This trend was far clearer in the root responses, particularly for the relationship between the Hg transcriptome response and RRG. In general, we expect more DEGs to be directly related to a greater stress response in less resilient genotypes (Bhat et al., 2024; Paape et al., 2016), but other unknown aspects not accounted for in our broad phenotyping measures of leaf and root responses likely accounts for some unpredictability due to the complexity of metal ion transport/homeostasis and different genetic backgrounds among the genotypes used in this study.

Using GO-enrichment, we were able to identify metal specific, genotype specific, and tissue specific responses among our DEGs. The most common shared GO-terms between the Cd and Hg treatments in both leaf and root tissues across all genotypes were cellular detoxification, response to oxidative stress, glutathione metabolism, and peroxidase activity, which are universal indicators of heavy metal stress in plants (Kumar and Trivedi, 2018; Moustakas, 2023). Consistent with the number of DEGs, the roots showed the strongest GO-enrichment signal, particularly for the low Cd tolerant genotype (HM075). The nature of the GO-terms for this genotype in roots were related to cell wall, cell division (e.g., DNA replication, microtubule-based movement), alcohol dehydrogenase activity, and deoxygenase activity, suggesting Cd stress in this genotype had large effects on fundamental biological processes in the roots of low tolerant genotype. While GO-enrichment is less granular than gene or gene family expression responses, it was largely consistent with patterns of expression of key gene families that were populated in the GO-terms such as UGTs genes, oxidative stress genes, glutathione S-transferase genes, and ABC-transporters. For Hg treated genotypes, the pattern clearly showed that the low Hg accumulator genotype (HM302) had the most GO-enrichment, and the most obvious genotype specific responses in these gene families. This suggests an elevated stress response to Hg stress in the low Hg accumulator. In contrast, the high Hg accumulating genotype (HM304) showed enrichment in GO-terms specifically linked to detoxification pathways, such as heme binding and glutathione transferase activity. This pattern suggests that high accumulators may employ a more selective and possibly alternative mechanism for Hg stress response, aiming to mitigate Hg toxicity more directly.

### 4.2 Heavy metal stress in the nodules of *M. truncatula* genotypes

The interaction between two different symbiotic rhizobia strains with two *M. truncatula* host plant genotypes resulted in significant differences in plant phenotypes and the transcriptome. The Hg-tolerant plant-rhizobium genotype combination HM304-AMp08 promoted shoot biomass and nodule number even in the absence of Hg stress. Despite the general inhibition of shoot, root, and nodule development in all host plant-rhizobia strain combination, the Hg accumulator plant (HM304) and the Hg-tolerant strain (AMp08) resulted in the least impact on shoot biomass. Together, our results indicate a significant genotype by genotype interaction between HM304 (high Hg accumulator) and AMp08 (high Hg tolerant rhizobia strain) which supports plant growth promotion by the AMp08 strain.

Because the high tolerant *S. medicae* strain AMp08 is known to have a Mer operon (Bhat et al., 2024), it can tolerate and evacuate high levels of Hg from cells in the form of Hg^0^. Moreover, the nitrogen fixation genes *nifA*, *nifT* and *nifB* were all significantly upregulated in the AMp08 strain while in symbiosis with the high accumulating *M. truncatula* genotype (HM304) (Bhat et al., 2024). The ability to eliminate Hg from the symbiosome of the nodule, in addition to upregulated *nif* expression may contribute to the higher nodule number and greater biomass traits with this host plant-rhizobia strain combination. We also note that a previous study that used AMp08 to inoculate *M. truncatula* host plants, showed no significant reduction in nitrogenase in plants treated with 500 µM HgCl_2_ compared with control plants (Arregui et al., 2021). Further support for the benefit of the symbiotic interaction with the high Hg tolerant strain conferred to the host plant was found in the XRF imaging which showed that the distribution of Fe was less impacted by Hg stress with the HT strain inoculated plants. It is well known that symbiosis, healthy nodules, and nitrogen fixation in legumes are highly Fe dependent (Escudero et al., 2020b, 2020a; González-Guerrero et al., 2016; Rodríguez-Haas et al., 2013). While the Mer operon is an attractive component to attribute plant growth promotion under Hg stress, a tight symbiosis from highly compatible partners can potentially provide enough resilience to stress due to optimized nitrogen fixation which strongly benefits host plants. We are therefore cautious to interpret the reduced impact of Hg stress in this plant-microbe combination exclusively to the Mer operon.

One of the most remarkable findings from the transcriptome, was the drastic reduction in the number of DEGs when both plant genotypes were inoculated with the high HT rhizobia strain compared with the LT strain. This strongly suggests that the resilient strain reduced the stress in the host plant by buffering the effects of Hg toxicity. Because strain quality and strong host-rhizobia compatibility is related to host plant fitness in legume-rhizobia symbiosis (Batstone et al., 2022; Mendoza-Suárez et al., 2020; Porter et al., 2011), the direct mechanism of a beneficial interaction as the result of the Hg tolerance of the *S. medicae* strain is not certain. However, the combination of reduced impact on the transcriptome combined with the reduced phenotypic effects when plants form symbiosis with the high HT strain suggests that rhizobia may perform a dual role in nitrogen fixation and also mediating toxic ion stress. In the present case, our results strongly support that the Mer operon is the most likely mediator of Hg stress due to its extremely efficient ability to transform inorganic Hg to a volatile form (Boyd and Barkay, 2012) that would be far less harmful to the host plant. It is also noteworthy that the same pattern was observed in rhizobia DEGs in nodules, whereby the HT strain had less than half the number of DEGs compared with the LT strain in response to Hg (Bhat et al., 2024). This suggests a unique property in the AMp08 strain that highlights the use of dual-transcriptomics to identify meaningful patterns of differential expression in both partners when genetic diversity is included in the experimental design.

The GO-enrichment clearly showed groups of non-overlapping gene ontologies with functional categories that were specific to plant genotype-rhizobia strain combinations which further illustrated genotype by genotype interactions that affected plant gene expression responses. The gene ontologies associated with the host plants inoculated with the low tolerant rhizobia strain showed disruption of fundamental biological processes such as cell division, and oxidative stress responses suggesting the nodule environment was highly toxic to the plant. When the high tolerant strain was used, particularly with the high Hg accumulating host plant, GO-terms often associated with heavy metal detoxification included ABC-transporter activity, heme/iron ion binding, glutathione transferase, UDP-glycosyltransferase activity, and several oxidative stress GO-terms. Among the GO-terms included with oxidative stress were flavonoid biosynthetic processes, and flavonoid pathway genes are known to be affected by Hg stress (Alvarez-Rivera et al., 2022) and flavonoids are known to trigger nod factor production in symbiotic bacteria, and are important throughout legume nodule development (Gifford et al., 2018). By contrast, the high Hg accumulating plant genotype in symbiosis with the low Hg tolerant rhizobia strain resulted in only oxidative stress response GO-term enrichment, and none of the ABC-transporter activity, heme/iron ion binding, glutathione transferase, or UDP-glycosyltransferase activity that were found in symbiosis with the high Hg tolerant strain. Moreover, none of the cell division or fundamental biological processes were enriched when in the high Hg accumulator plant-high Hg tolerant rhizobia combination.

Consistent with the genome wide data, the subset of symbiosis genes also showed the same trend of DEGs when the HT strain was used. The pattern of expression of this set of genes stood out most strongly in the high Hg accumulating plant genotype (HM304) and HT strain (AMp08) combination (Figure 7c, e), where the small number of plant symbiosis genes appeared least affected by the Hg stress. While the functions of this subset of genes varies widely in the process of signaling, nodulation, nitrogen fixation, and symbiosis as a whole, the patterns (and direction, whether up or down-regulated) of expression in combination with the plant phenotypes before and after Hg stress, suggests that the metal resilience in the host plant and rhizobia partners likely assist in mediated toxic ion stress. This specific combination of host plant and rhizobia strain also indicates a genotype by genotype affect that could be optimized for greater resilience to contaminated environments and for bioremediation (Fagorzi et al., 2018; Naguib et al., 2018; Narayanan and Ma, 2023; Siripornadulsil and Siripornadulsil, 2013). While our example of a putative detoxification mechanism by a genetically adapted rhizobia strain is unique to Hg in the present case, there is need to further explore mechanisms that employ rhizobia to contribute to retention and compartmentalization in nodules of legumes, as the majority of research has focused on micronutrients in nodules (González-Guerrero et al., 2023). If toxic metals could be contained in the edible parts of the host plant, there are benefits to using adapted plant-microbe interactions that extend beyond bioremediation.

## Acknowledgements

Funding for this project came from the Department of Energy Quantitative Plant Science Initiative SFA, USDA-ARS Project Number 3092-53000-001-000D, and the Agencia Estatal de Investigación, Spain, grant number PID2021-125371OB-I00 awarded to JJP. Sequencing of *M. truncatula* seedlings in hydroponics was done with a Community Science Project (proposal: 10.46936/10.25585/60001308) awarded to TP, conducted by the U.S. Department of Energy Joint Genome Institute (https://ror.org/04xm1d337), a DOE Office of Science User Facility, is supported by the Office of Science of the U.S. Department of Energy operated under Contract No. DE-AC02-05CH11231. We thank Nevin Young and Roxanne Denny at the University of Minnesota for germplasm, and their support and discussions regarding the Medicago HapMap collection. We thank Jeremy Schmutz, Kerrie Berry and T.B.K Reddy from JGI for support with our Community Science Project and data management. We thank Tiffany Victor and Ryan Tappero at NSLS-II, Brookhaven National Laboratory for training and support with X-Ray fluorescence microscopy. We would like to thank Katy Heath, Alvaro Hernandez and Chris Wight at the University of Illinois at Urbana-Champaign for consulting, oligo design, and rRNA cleanup and sequencing for the nodule samples. The authors declare that they have no competing interests.

## Tables

See Supplementary Tables and Figures document.

## Data Accessibility

Expression data for whole genome the whole and lists of differentially expression genes can be found in Supplementary Data File 1.

Input data for main figures and supplementary figures can be found in Supplementary Data File 2.

Lists of differentially expressed symbiosis related genes from nodules corresponding to each plant-rhizobia genotype can be found in Supplementary Data File 3.

Raw reads from *Medicago truncatula* hydroponics experiment (150 bp paired-end Illumina reads). https://www.ncbi.nlm.nih.gov/bioproject/?term=PRJNA994975

Raw reads from dual-transcriptomics (150 bp paired-end Illumina reads). https://www.ncbi.nlm.nih.gov/bioproject/?term=PRJNA986885

## Notes

### Competing Interest Statement

The authors have declared no competing interest.

