## SupplementaryMaterial for "Plant genotype and microbial strain combinations strongly influence the transcriptome under heavy metal stress conditions"

### Supplementary Figures.

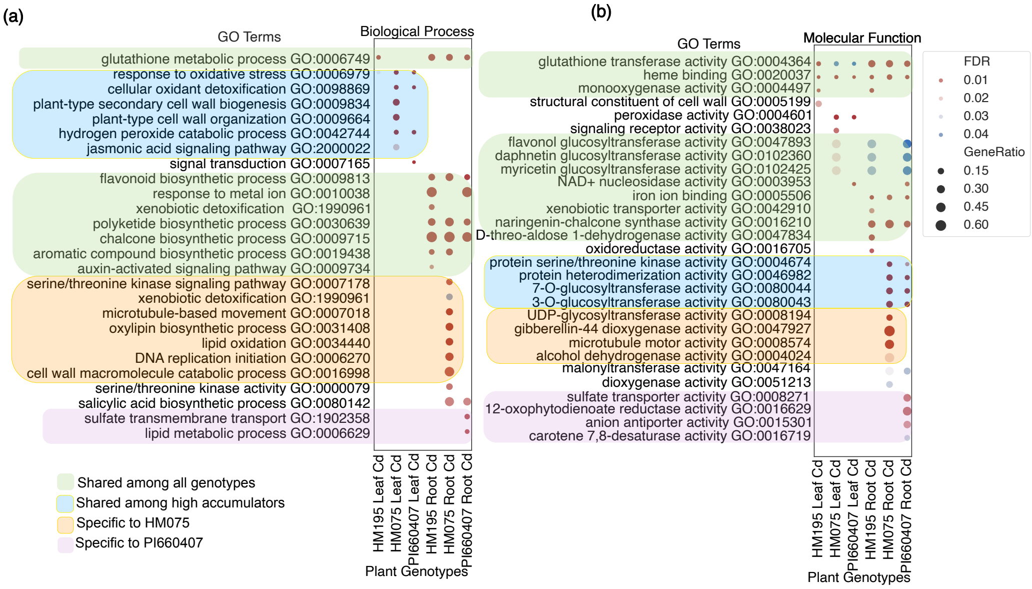


Fig. S1. Gene ontology (GO) enrichment analysis of DEGs in leaf and root tissues of *M. truncatula* genotypes showing the Cd treated genotypes HM195 (low accumulation), HM075 (high accumulation) and PI660407 (high tolerance and high accumulation). The dot-plot shows enriched GO-terms for DEGs using biological processes (a) and molecular functions (b), where the y-axis is the name of GO-terms along with their GO IDs. DEGs used for GO-enrichment were defined as $log_2FC \geq 1$ and adjusted p-value $< 0.05$. The x-axes of (a) and (b) are the plant genotypes, metal treatment and tissue (leaf or root). The color of the circle indicates the false discovery rate (FDR) adjusted p-value of over-enriched GO-terms. The gene ratio is the percentage of total DEGs in the given GO-term and circle size corresponds to relative abundance of genes in that GO term. We grouped sets of GO-terms with light green shaded areas corresponding to shared GO-terms among all genotypes, blue shaded for GO-terms associated with high Cd accumulator genotypes, light blue for GO-terms specific to HM075 and light pink for GO-terms specific to PI660407 genotype.

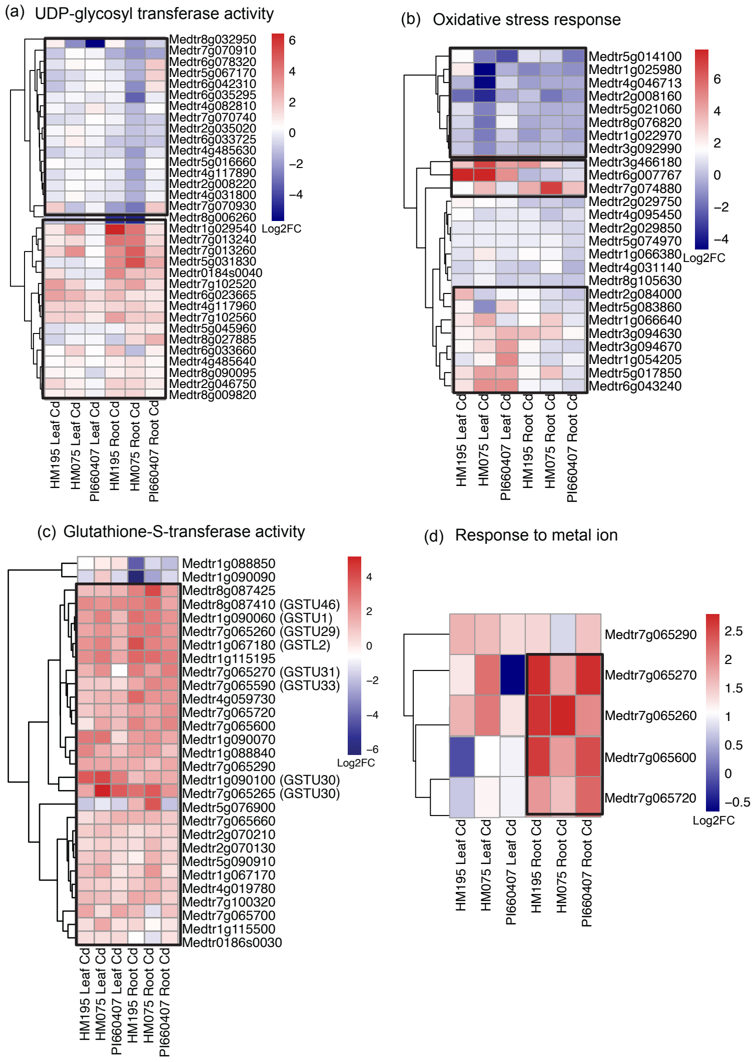


Fig. S2. Heatmaps showing genes expressed among Cd treated genotypes within the GO-terms from Fig. 2; (a) UDP-glycosyl transferase (b) oxidative stress response (c) glutathione-s-transferase and (d) response to metal ion. The heatmap rows (y-axis) represent gene IDs with gene family names, and the columns (x-axis) correspond to *M. truncatula* plant genotypes, metal treatments, and tissues (leaf or root). Each cell is colored based on the $log_2FC$ expression in treated conditions compared to control conditions of each gene labeled on y-axis. The scale bar corresponds to upregulation shown in red and downregulation shown in blue using the $log_2FC$ scale. The black boxes include the gene clusters with variable expression patterns among genotypes. Clustering of expression is indicated by the dendrogram on the left y-axis, while the x-axis samples were constrained to remain in the same order for leaf and root.

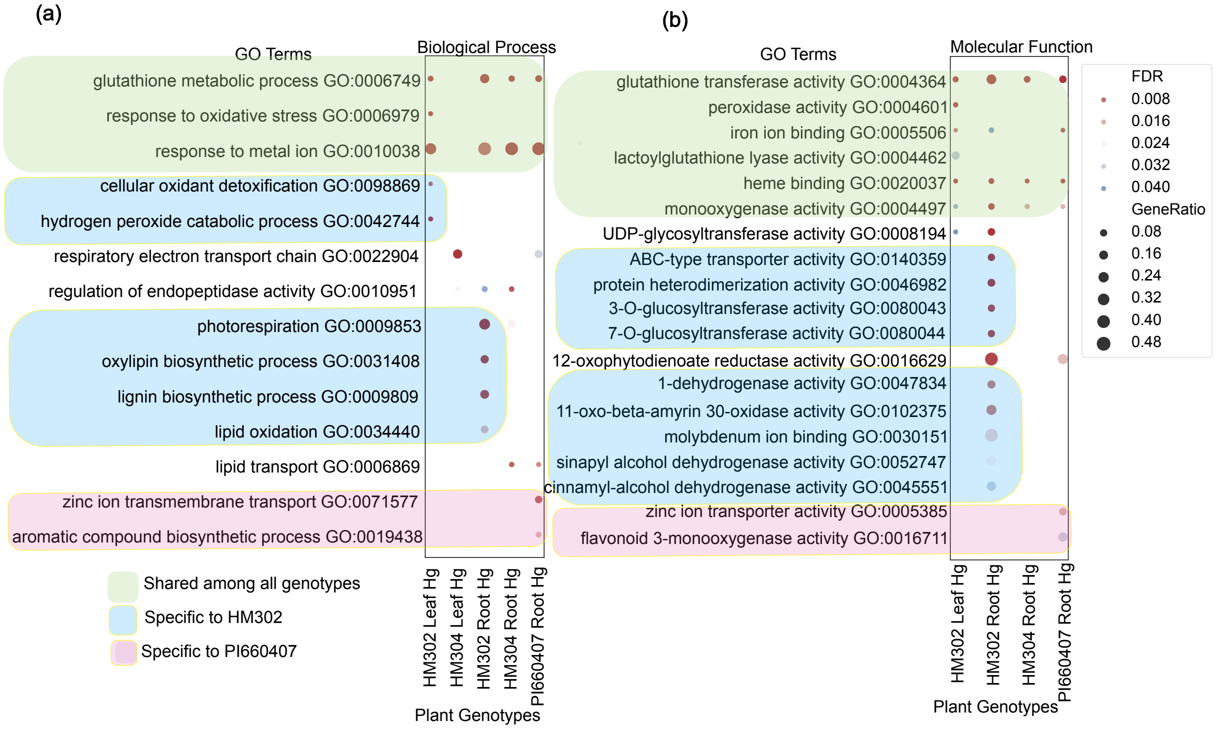


Fig. S3. Gene ontology (GO) enrichment analysis of DEGs in leaf and root tissues of the Hg-treated *M. truncatula* genotypes HM302 (low accumulation), HM304 (high accumulation) and PI660407 (high accumulation and high tolerance). The dot-plot shows enriched GO-terms for DEGs using biological processes (a) and molecular functions (b), where the y-axis is the name of GO-terms along with their GO IDs. DEGs used for GO-enrichment were defined as $\log_2 FC \geq 1$ and adjusted p-value < 0.05. The x-axes of (a) and (b) are the plant genotypes, metal treatment and tissue (leaf or root). The color of the circle indicates the false discovery rate (FDR) adjusted p-value of over-enriched GO-terms. The gene ratio is the percentage of total DEGs in the given GO-term and circle size corresponds to relative abundance of genes in that GO term. We grouped sets of GO-terms with light green shaded areas corresponding to shared GO-terms among all genotypes, blue shaded for GO-terms specific to HM302 and light pink for GO-terms specific to PI660407 genotype.

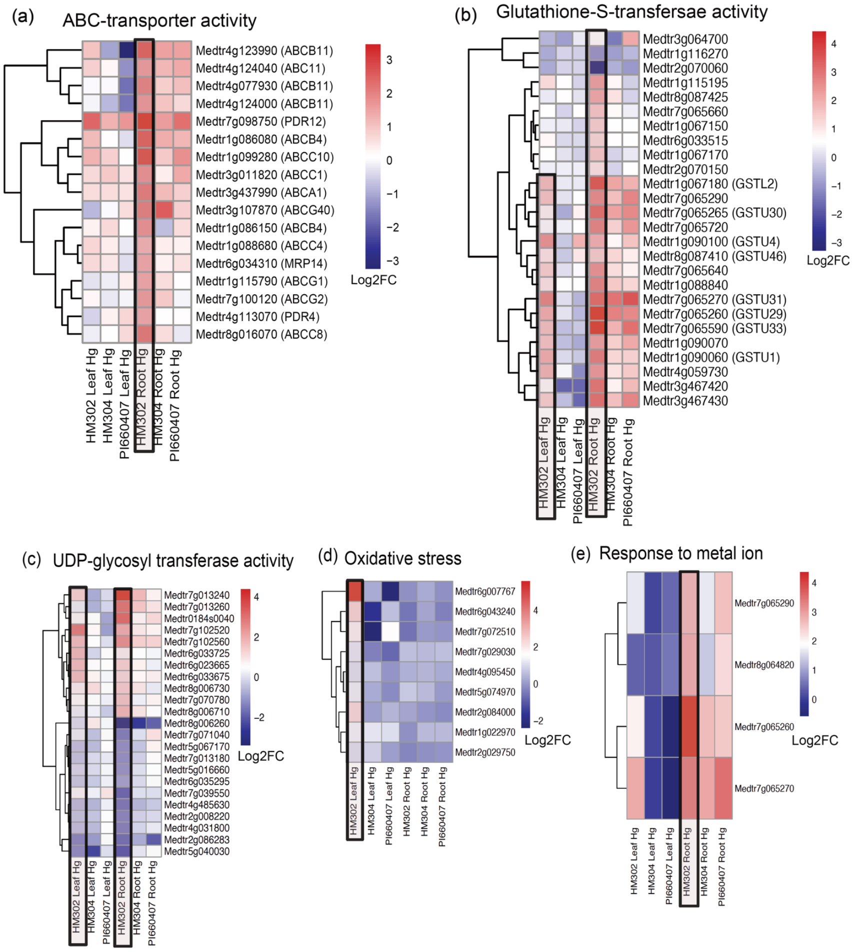


Fig. S4. Heatmaps showing genes expressed among Hg treated genotypes within the GO-terms from Fig. 2 and Fig. S2; (a) ABC-transporter activity, (b) glutathione-s-transferase, (c) UDP-glycosyl transferase (d) oxidative stress, and (e) response to metal ion. The heatmap rows (y-axis) represent gene IDs with gene family names, and the columns (x-axis) correspond to *M. truncatula* plant genotypes, metal treatments, and tissues (leaf or root). Each cell is colored based on the $\log_2$FC expression in treated conditions compared to control conditions of each gene labeled on y-axis. The scale bar corresponds to upregulation shown in red and downregulation shown in blue using the $\log_2$FC scale. The black boxes include the gene clusters with variable expression patterns among genotypes. Clustering of expression is indicated by the dendrogram on the left y-axis, while the x-axis samples were constrained to remain in the same order for leaf and roots.

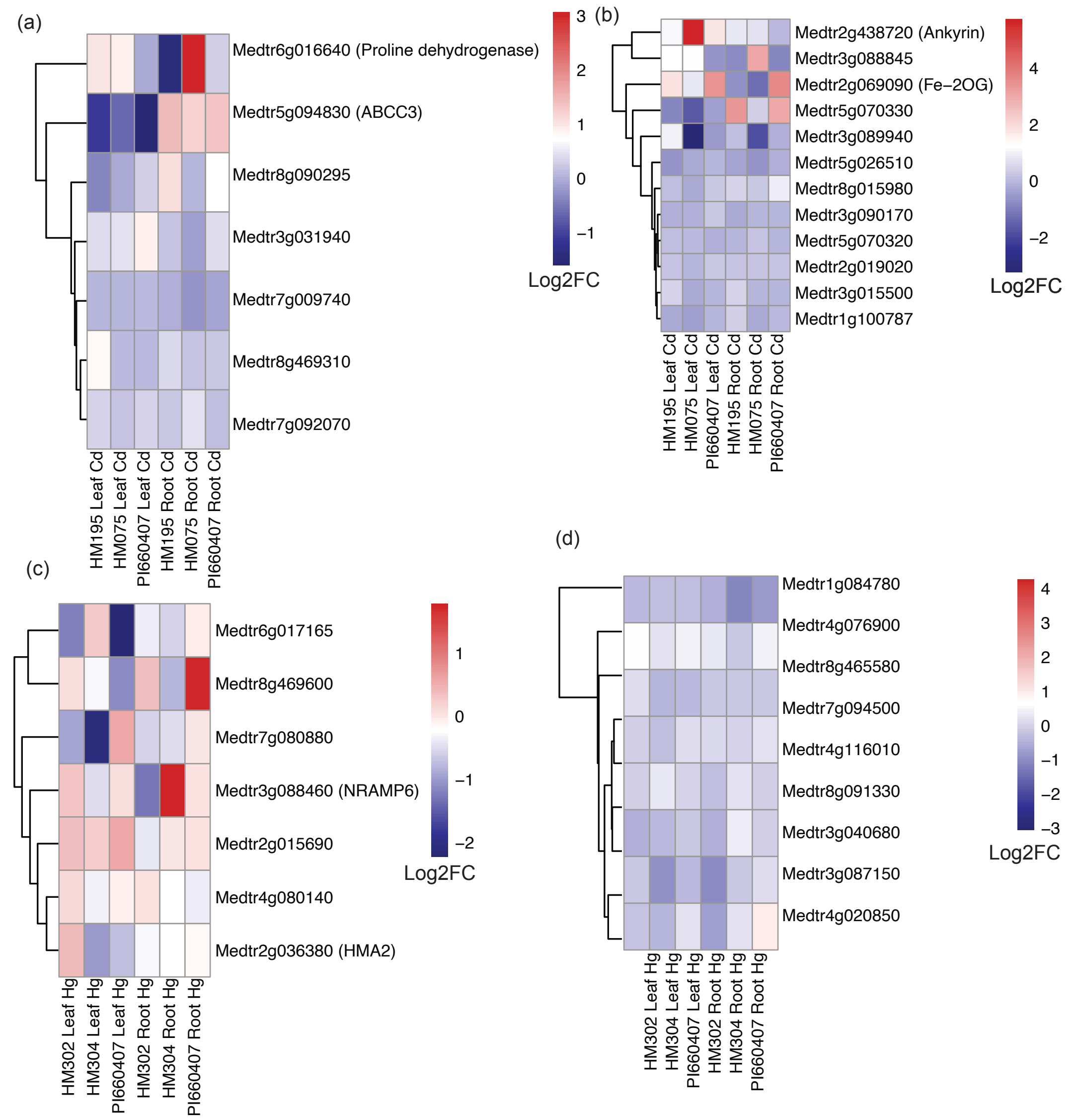


Fig. S5. Heatmaps showing the genes found common among our transcriptomic data and GWAS study in (Paape et al., 2022) which corresponds to phenotypic traits of *M. truncatula* genotypes under Cd and Hg stress i.e., metal accumulation (from leaf tissue) and relative root growth (RRG) where (a) Cd accumulation, (b) Cd RRG, (c) Hg leaf and (d) Hg RRG. The heatmap rows (y-axis) represent gene IDs with gene family names, and the columns (x-axis) correspond to *M. truncatula* plant genotypes, metal treatments, and tissues (leaf or root). Each cell is colored based on the expression level of the respective gene in treated conditions compared to control conditions measured as $\log_2$FC. The scale bar corresponds to upregulation shown in red and downregulation shown in blue using the $\log_2$FC scale. Clustering of expression is indicated by the dendrogram on the left y-axis, while the x-axis samples were constrained to remain in the same order for leaf and roots.

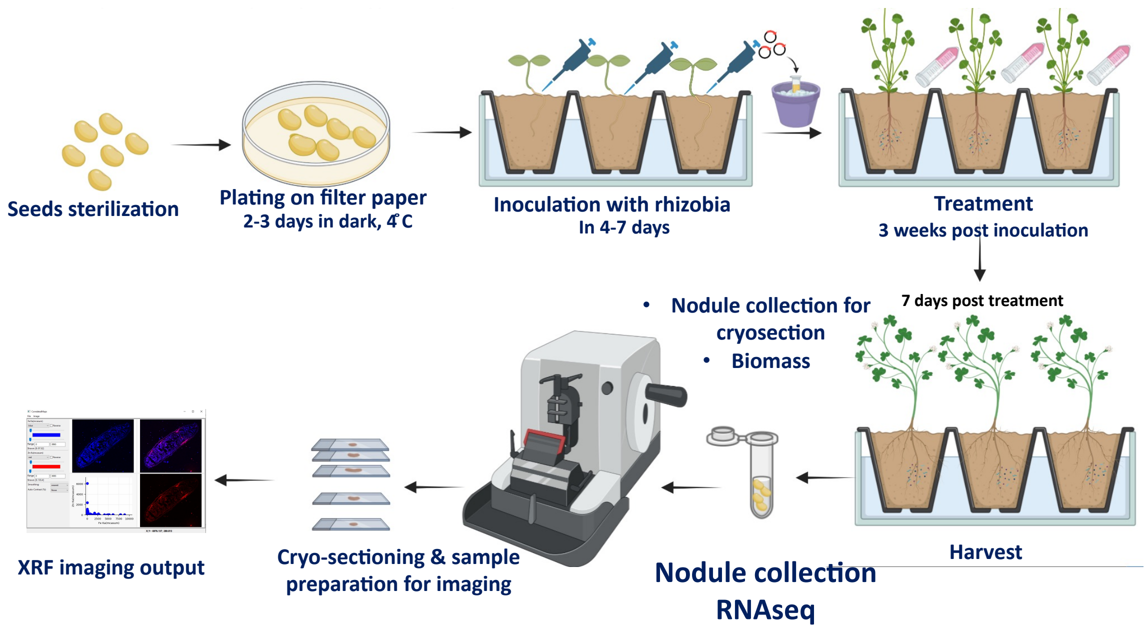


Fig. S6. Experimental design for plant growth, metal treatment and sample collection for phenotyping, dual-transcriptomics, and XRF-imaging.

(a)

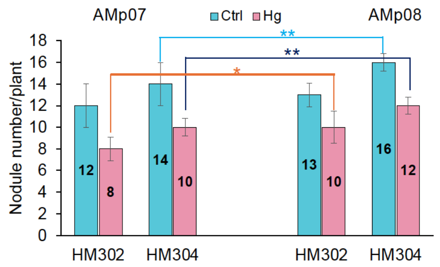


(b)

Statistical analysis among host genotype and rhizobia strain combinations under control and Hg treatment

| | p-Value |
|---|---|
| **Ctrl** | |
| HM302 AMp08/HM302 AMp07 | 0.243021 |
| HM304 AMp08/HM304/AMp07 | 0.00404 |
| **Hg** | |
| HM302 AMp08/HM302 AMp07 | 0.023012 |
| HM304 AMp08/HM304/AMp07 | 0.000199 |

(c) (d)

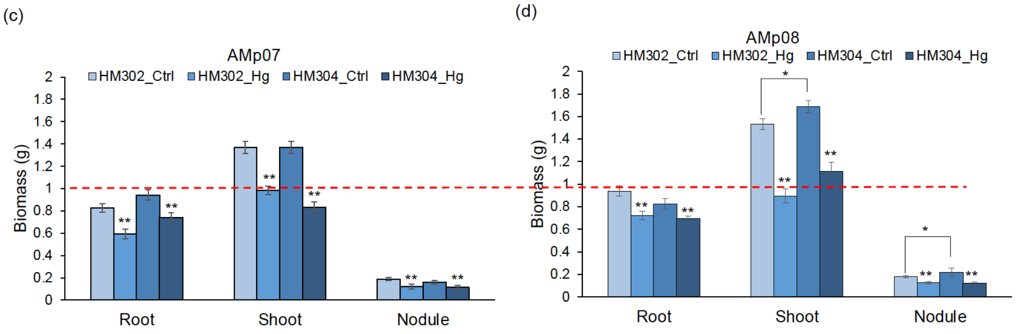


Fig. S7. Nodule number and biomass yield of root, shoot, and nodules were measured in two *M. truncatula* genotypes: HM302 (Low Hg-accumulator) and HM304 (High Hg-accumulator), inoculated with *S. medicae* strains AMp07 (LT) or AMp08 (HT) rhizobia strains. (a) Bar plot showing the average number of nodules per plant after inoculation with AMp07 and AMp08 in control (Ctrl) and Hg-treated (Hg) conditions. These measurements included five biological replicates and error bars are ± SD from the mean. The orange line aids in comparing the HM302-AMp07 and HM302-AMp08 under Hg stress, dark blue line shows comparison among HM304-AMp07 and HM304-AMp08 under Hg stress and light blue line shows comparison among AMp07 and AMp08-inoculated HM304 genotype under control conditions. (b) Statistical analysis among host genotype and rhizobia strain combination under control and Hg treatment from comparisons made in (a). The biomass measurements of fresh roots, shoots, and nodules in both non-treated and Hg-treated is depicted in Fig.s (c) both plant genotypes inoculated with AMp07 and (d) both plant genotypes inoculated with AMp08. In each graph, the y-axes represent biomass (in grams), while the x-axes indicate different tissue types. The dashed red was placed at the value of one for the comparison between the AMp07 and AMp08-inoculated plants. The data represent the means ± SD from 5 independent replicates. Statistical tests were performed with two-sided student *t*-test related to non-treated samples. (**p < 0.01, *<0.05).

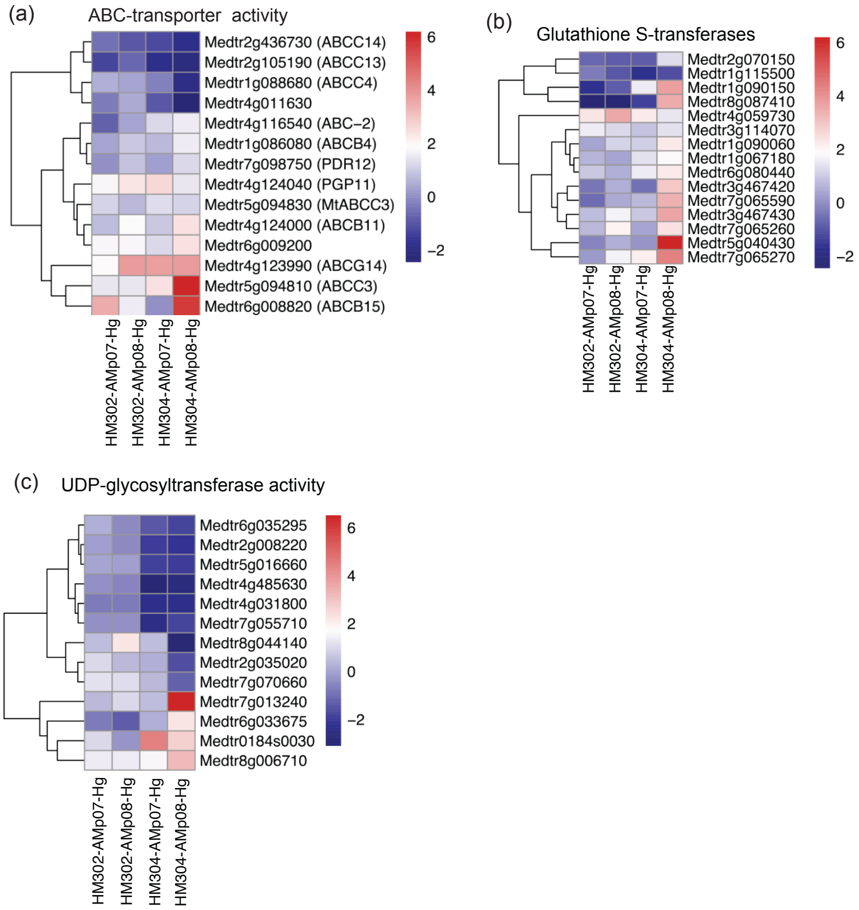


Fig. S8. Heatmaps showing genes in the GO-term, ABC-transporter activity (a), Glutathione-S-transferase (b) response to oxidative stress (c) and UDP-glycosyl transferase associated with Hg stress response in Fig. 6. The heatmap rows (y-axis) represent gene IDs with gene family names, and the columns (x-axis) correspond to *M. truncatula* plant genotypes, metal treatments, and tissues (nodules). Each cell is colored based on the $\log_2$FC expression in treated conditions compared to control conditions of each gene labeled on y-axis. The scale bar corresponds to upregulation shown in red and downregulation shown in blue using the $\log_2$FC scale. Clustering of expression is indicated by the dendrogram on the left y-axis.

**Supplementary Tables.**

Table S1. Selected *Medicago truncatula* genotypes and their phenotypic traits under Cd and Hg treatment when grown under hydroponic conditions.

| Genotype | Treatment | Tolerance Level | Accumulation Level |
|---|---|---|---|
| HM195 | Cd | Low | Low |
| HM075 | Cd | Low | High |
| PI660407 | Cd | High | High |
| HM302 | Hg | Low | Low |
| HM304 | Hg | Low | High |
| PI660407 | Hg | High | Low |

Table S2. Differentially Expressed Genes (DEGs) in leaf and root tissues of *Medicago truncatula* genotypes under Cd and Hg treatment (total DEGs, Upregulated DEGs and Downregulated DEGs).

| Genotype | Tissue | Treatment | Total DEGs | Upregulated DEGs | Downregulated DEGs |
|---|---|---|---|---|---|
| HM195 | Leaf | Cd | 547 | 372 | 175 |
| HM195 | Root | Cd | 1587 | 1047 | 540 |
| HM075 | Leaf | Cd | 1351 | 884 | 467 |
| HM075 | Root | Cd | 4018 | 2312 | 1706 |
| PI660407 | Leaf | Cd | 382 | 281 | 101 |
| PI660407 | Root | Cd | 1566 | 1210 | 356 |
| HM302 | Leaf | Hg | 278 | 189 | 89 |
| HM302 | Root | Hg | 1274 | 739 | 535 |
| HM304 | Leaf | Hg | 250 | 93 | 157 |
| HM304 | Root | Hg | 585 | 346 | 239 |
| PI660407 | Leaf | Hg | 30 | 2 | 28 |
| PI660407 | Root | Hg | 348 | 293 | 55 |
| HM302-AMp07 | Nodules | Hg | 3429 | 1723 | 1706 |
| HM302-AMp08 | Nodules | Hg | 1899 | 1131 | 768 |
| HM304-AMp07 | Nodules | Hg | 3643 | 2191 | 1452 |
| HM304-AMp08 | Nodules | Hg | 1143 | 775 | 368 |

Table S3-S6. The tables next to the volcano plots show the genes and functional annotations from Fig. expression levels, and statistical significance. These tables serve as a visual representation, elucidating the differential expression patterns and the significance of gene expression changes across the genotypes.

Table S3

HM302_AMp07

| Symbol | Gene | Function | Log2FC | padj |
|---|---|---|---|---|
| MtrunA17_Chr3g0135351 | Medtr3g103960 | Zinc finger family protein | -3.87 | 8.58E-19 |
| MtrunA17_Chr3g0102071 | Medtr3g058630 | Zinc transporter 11 precursor | -3.08 | 6.36E-13 |
| MtrunA17_Chr6g0451911 | Medtr6g008820 | ABC transporter family protein | 3.46 | 1.69E-12 |
| MtrunA17_Chr6g0487091 | Medtr6g088805 | MATE efflux family protein | 4.7 | 1.73E-11 |
| MtrunA17_Chr1g0176441 | Medtr1g054935 | ABC-2 type transporter family protein | -1.69 | 5.60E-11 |
| MtrunA17_Chr7g0243711 | Medtr7g070740 | UDP-glucosyl transferase | 2.84 | 3.69E-09 |

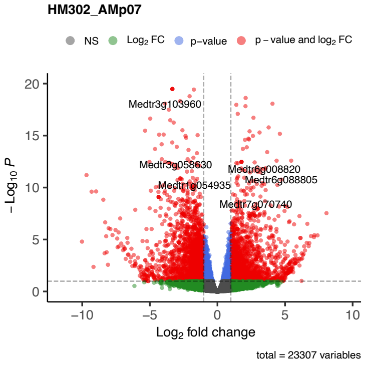


Table S4

HM302_AMp08

| Symbol | Gene | Function | Log2FC | padj |
|---|---|---|---|---|
| MtrunA17_Chr4g0069351 | Medtr4g123990 | P-glycoprotein 11 | 3.83 | 1.53E-28 |
| MtrunA17_Chr5g0409121 | Medtr5g026620 | Alternative oxidase 1A | 2.74 | 2.07E-26 |
| MtrunA17_Chr4g0029341 | Medtr4g059730 | Glutathione S-transferase TAU 8 | 3.45 | 1.02E-21 |
| MtrunA17_Chr4g0046331 | Medtr4g088170 | Cytochrome P450 | 3.09 | 2.99E-19 |
| MtrunA17_Chr2g0324591 | Medtr2g090525 | Proton pump interactor 1 | 8.92 | 9.78E-08 |
| MtrunA17_Chr5g0426431 | Medtr5g464810 | UDP-glucosyl transferase 85A2 | -1.31 | 5.47E-06 |
| MtrunA17_Chr7g0217461 | Medtr7g010820 | Nitrate transporter 1.5 | -4.44 | 1.22E-05 |
| MtrunA17_Chr1g0175561 | Medtr1g054205 | Peroxidase 2 | 7.43 | 9.18E-05 |

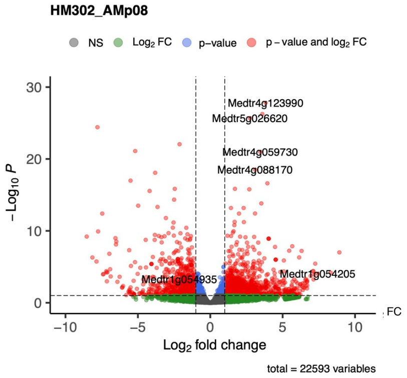


Table S5

HM304_AMp07

| Symbol | Gene | Function | Log2FC | padj |
|---|---|---|---|---|
| MtrunA17_Chr7g0243241 | Medtr7g069980 | Ferretin 1 | -8.20 | 3.05E-32 |
| MtrunA17_Chr8g0388921 | Medtr8g100135 | cytochrome P450, family 716 | 3.02 | 5.59E-39 |
| MtrunA17_Chr5g0426431 | Medtr5g464810 | UDP-glucosyl transferase 85A2 | 3.25 | 9.03E-20 |
| MtrunA17_Chr4g0069351 | Medtr4g123990 | P-glycoprotein 11 | 3.71 | 3.46E-30 |
| MtrunA17_Chr4g0050811 | Medtr4g094325 | Vacuolar iron transporter (VIT) family protein | 3.72 | 1.23E-11 |
| MtrunA17_Chr6g0487091 | Medtr6g088805 | MATE efflux family protein | 6.05 | 0.00096883 |
| MtrunA17_Chr7g0244461 | Medtr7g072510 | Peroxidase superfamily protein | 7.87 | 3.68E-08 |

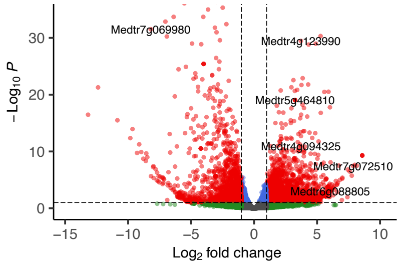


Table S6

HM304_AMp08

| Symbol | Gene | Function | Log2FC | padj |
|---|---|---|---|---|
| MtrunA17_Chr1g0189841 | Medtr1g079490 | UDP_glycosyl transferase | 3.12 | 1.88E-31 |
| MtrunA17_Chr2g0296671 | Medtr2g437910 | Vacuolar sorting receptor | 2.32 | 4.55E-17 |
| MtrunA17_Chr6g0453201 | Medtr6g012170 | Oxidoreductase, Zinc-binding dehydrogenase protein | -3.17 | 2.33E-14 |
| MtrunA17_Chr3g0107741 | Medtr3g467430 | Glutathione S-transferase TAU 20 | 3.52 | 4.54E-12 |
| MtrunA17_Chr7g0239111 | Medtr7g062580 | Fe(II)-dependent oxygenase protein | 3.14 | 2.15E-08 |
| MtrunA17_Chr2g0331081 | Medtr2g101090 | ABC-2 and PDR-ABC-type transporter | 3.24 | 5.80E-07 |

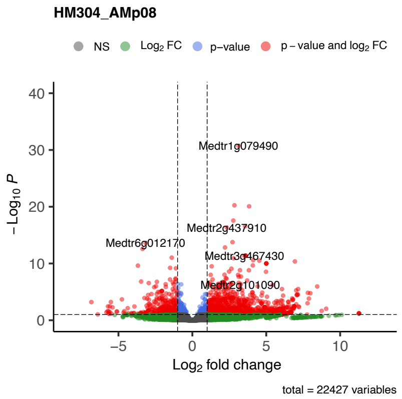
